# Neanderthal introgression reintroduced functional ancestral alleles lost in Eurasian populations

**DOI:** 10.1101/533257

**Authors:** David C. Rinker, Corinne N. Simonti, Evonne McArthur, Douglas Shaw, Emily Hodges, John A. Capra

## Abstract

Neanderthal ancestry remains across modern Eurasian genomes, and introgressed sequences influence diverse phenotypes, including immune, skin, and neuropsychiatric diseases. Interpretation of introgressed sequences has focused on alleles derived in the Neanderthal lineage. Here, we demonstrate that introgressed Neanderthal haplotypes also carry hundreds of thousands of ancestral alleles that had been lost in Eurasian populations. These reintroduced alleles (RAs) exist exclusively on Neanderthal haplotypes in Eurasian populations. Illustrating the broad potential influence of these RAs, we find that over 70% of known phenotype associations with introgressed Neanderthal-derived alleles (NDAs) are equally associated with RAs. We also discover enrichment for RAs among introgressed eQTL in many tissues, including more than half of the brain tissues analyzed. Finally, combining expression quantitative trait loci (eQTL), massively parallel reporter assay (MPRA) data, and *in vitro* validation, we show that RAs can regulate gene expression independent of NDAs. In summary, our study reveals that Neanderthal introgression supplied Eurasians with many lost ancestral variants that may have restored lost regulatory functions. Thus, RAs should be considered when evaluating the effects of introgression.

**ONE SENTENCE SUMMARY:** Neanderthal interbreeding with anatomically modern humans restored thousands of ancient alleles that were previously lost in Eurasian populations.

## MAIN TEXT

Modern Eurasian populations have significantly lower genetic diversity than modern African populations, despite having larger census population sizes (*1, 2*). This disparity reflects the genetic bottlenecks experienced by the direct ancestors of Eurasian anatomically modern humans (AMH) as they moved out of Africa approximately 50,000 years ago (*2, 3*). The effective population size of this ancestral Eurasian population is estimated to have been less than 20% of the size of the contemporaneous African populations (*1, 4*). As a result of this out of Africa (OOA) bottleneck and subsequent population dynamics, millions of ancient alleles were lost in the ancestors of modern Eurasian populations.

More than 500,000 years prior to the Eurasian OOA bottleneck, members of other hominin groups in Africa, including the ancestors of Neanderthals and Denisovans, moved into Eurasia (*5*). The sequencing of ancient DNA from Neanderthal and Denisovan individuals has enabled reconstruction of their genomes (*5*–*7*). Comparing Neanderthal genomes to genomes of modern humans from around the world revealed that Eurasian AMHs interbred with Neanderthals approximately 50,000 years ago (*5, 8*). The legacy of this archaic introgression is reflected in the genomes of modern Eurasians, where ∼1–3% of individuals’ DNA sequences are of Neanderthal ancestry (*9*–*12*).

Neanderthal introgression introduced many new alleles into Eurasian populations that were derived on the Neanderthal lineage. It has been hypothesized that some of these alleles were adapted to non-African environments and thus were beneficial to Eurasian AMH (*9, 10, 13*–*18*). However, Neanderthal interbreeding also likely came with a genetic cost due to accumulation of weakly deleterious alleles in their lineage, because of their lower effective population size compared to AMHs (*19, 20*). Indeed, the distribution of archaic ancestry across modern Eurasian genomes is non-random, with significant deserts of Neanderthal ancestry as well as many genomic regions in which Neanderthal ancestry is common. This distribution is generally attributed to the long term effects of positive and negative selection acting on introgressed Neanderthal alleles (*9, 10, 21*), with negative selection acting most strongly immediately after admixture (*22*).

Introgressed alleles on Neanderthal haplotypes that remain in modern Eurasian populations are associated with diverse traits, including risk for skin, immune, and neuropsychiatric diseases (*13, 14, 23*–*26*). For example, an introgressed Neanderthal haplotype at the *OAS1* locus is associated with innate immune response; however, this haplotype also contains an ancient hominin allele in high linkage disequilibrium (LD) with the Neanderthal alleles that could influence function (*27*). Thus, while most studies have focused on identifying and testing the effects of Neanderthal derived alleles in AMHs, archaic admixture may also have served as a route by which more ancient functional alleles reentered the genomes of Eurasians (*27, 28*).

Here, we explore the hypothesis that Neanderthal introgression reintroduced previously lost ancestral alleles into Eurasian populations. To evaluate this hypothesis, we analyze archaic, modern, and simulated genomes to characterize the prevalence of the reintroduction of lost alleles to Eurasians. Given the conservation of many of these alleles in Africans and in closely related ape species, we evaluate and test whether the reintroduction of some of these variants may also have restored lost functions. We identify more than 200,000 ancient alleles that are only present on introgressed Neanderthal haplotypes in modern Eurasian populations. We discover enrichment for reintroduced alleles among introgressed haplotypes with gene regulatory effects in several tissues, including the brain. We demonstrate functional effects for reintroduced alleles using computational analyses, cross-population comparisons of eQTL, and MPRA data. Finally, we experimentally validate the gene regulatory effects of a reintroduced allele independent of associated Neanderthal alleles in the context of both African and Eurasian haplotypes. Taken together, our results demonstrate that Neanderthal populations served as reservoirs of functional ancestral alleles that were lost to the ancestors of Eurasians (and in some cases all modern humans), and that some of these alleles have functional effects in Eurasians after being reintroduced by Neanderthal admixture.

## RESULTS

To illustrate the evolutionary scenarios we investigate here, consider the simple model of recent hominin demography presented in **Figure** 1A. Many alleles segregating in ancestral hominins were lost in the AMH lineage after the divergence of the ancestors of AMHs and Neanderthals. Some were lost in all AMHs, while others were lost only in Eurasian populations, e.g., during the OOA bottleneck. These lost alleles thus had the potential to be reintroduced into Eurasian populations via archaic admixture. Within these populations, reintroduced alleles would initially only be present on introgressed Neanderthal haplotypes, and over time many would retain high LD with Neanderthal-derived alleles in modern Eurasians. In the following analysis, we will refer to alleles that were present in the most recent common ancestor of AMHs and Neanderthals as “ancestral hominin alleles.”

**Figure 1.**
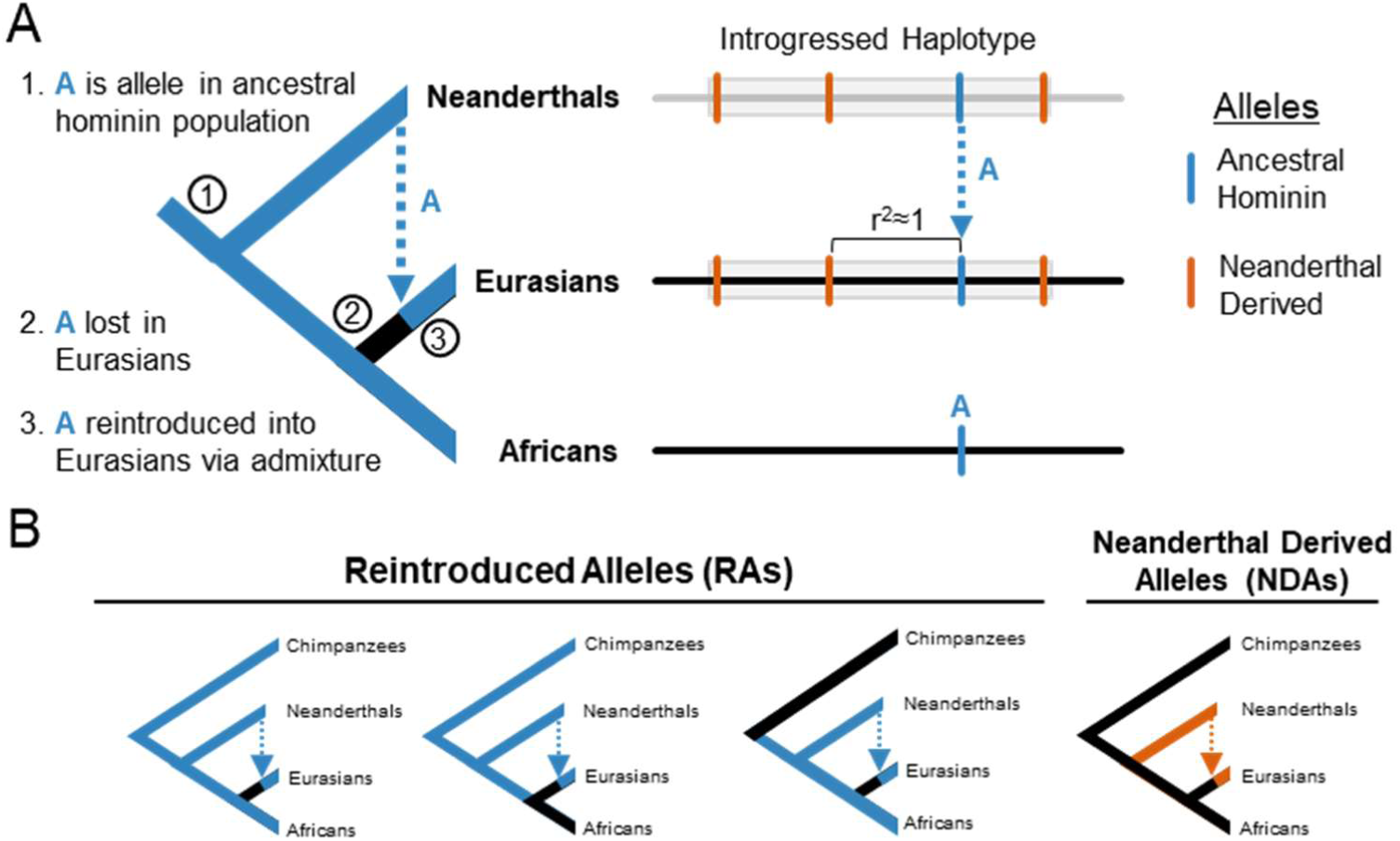
Schematic of the reintroduction of lost ancestral alleles by Neanderthal introgression. (A) Illustration of the evolutionary trajectory and resulting genomic signature of an allele A (blue) that was: (1) segregating in the ancestors of anatomically modern humans (AMHs) and Neanderthals, (2) lost to the ancestors of Eurasians in the human out of Africa (OOA) bottleneck or other subsequent migrations, and (3) reintroduced to Eurasians through Neanderthal admixture. Consequently, reintroduced alleles (RAs) are expected to be in high linkage disequilibrium with some Neanderthal-derived alleles (NDAs; orange) on introgressed haplotypes (gray) in modern Eurasians. (B) Schematics of the different evolutionary histories of interest in this paper. Alleles lost in Eurasians (or all AMHs) and reintroduced by Neanderthal introgression are referred to as RAs. Alleles that appeared in the Neanderthal lineage, were not present in the ancestors of humans and Neanderthals, and only exist on introgressed haplotypes in modern humans are referred to as Neanderthal-derived alleles (NDAs).

We will refer to ancestral hominin alleles that are only observed in Eurasians on introgressed Neanderthal haplotypes as reintroduced alleles (RAs), and introgressed alleles that first appeared on the Neanderthal lineage as Neanderthal-derived alleles (NDAs) (**Figure** 1B). Here we evaluate the presence and function of RAs in modern Eurasians and contrast them with NDAs.

### Hundreds of thousands of RAs exist in modern Eurasian populations

To identify candidate RAs in the genomes of modern Eurasians, we sought variants in 1000 Genomes Phase 3 Eurasian populations that are present only on introgressed haplotypes (**Figure** S3, **Methods**). We began with sets of previously identified SNPs that tag introgressed haplotypes in Eurasians. These SNPs were identified by S* and comparisons between Neanderthal genomes and the genomes of European (EUR), East Asian (EAS), and South Asian (SAS) populations (*12*). For each population, we identified candidate RAs by collecting variants that are in perfect LD (r^2^=1) with a Neanderthal tag SNP, but that are not tag SNPs themselves. We then evaluated each of these candidate RAs with regard to its ancestral status and presence in modern sub-Saharan Africans. Candidate alleles that matched the high-confidence ancestral allele call from 1000 Genomes or that were present at a frequency of >1% in sub-Saharan African populations without substantial Neanderthal ancestry were deemed RAs. Overall, ∼73% of Neanderthal tag SNPs are in perfect LD with at least one classifiable RA. Forward-time evolutionary simulations suggest that false positives due to recombination artefacts or convergent mutations, even at highly mutable CpG dinucleotides, are rare (**Figure** S1, S2; **Table** S1). Some Neanderthal haplotypes are present in some sub-Saharan African populations due to backflow post-admixture from Eurasians; however, given that the resulting fraction of Neanderthal ancestry is estimated to be very low (0.18%) (*6, 29*), our classification criteria prevent them from leading to many false positives (Methods). Finally, our approach is likely conservative, because many true RAs are not expected to retain perfect LD with any NDA.

Altogether, we identified 209,176 RAs (**Figure** 2B, **Figure** S3). The South Asian and East Asian populations each have more RAs (139,270 and 125,257, respectively) than the European populations (90,121). These numbers likely reflect the differences in the number of Neanderthal tag SNPs found in each population (**Figure** S3, **Figure** S4), the greater levels of Neanderthal ancestry previously observed in East Asians (*30, 31*), and the differences in the demographic history of these populations (*32*).

**Figure 2.**
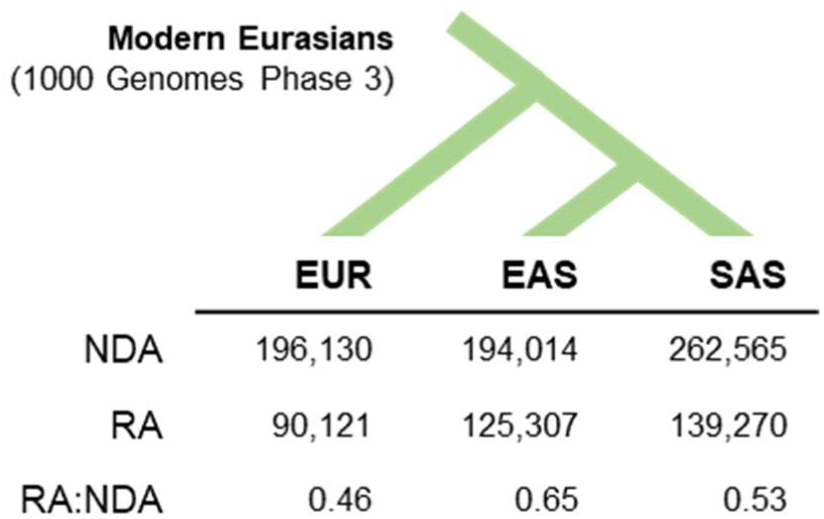
Neanderthal introgression reintroduced thousands of lost ancestral alleles to Eurasian populations. The number of RAs and NDAs in each Eurasian 1000 Genomes population (EAS = East Asian; EUR = European ancestry; SAS = South Asian) identified by our pipeline (Figure S3; Methods). Overall, Neanderthal admixture is responsible for the presence of over 200,000 ancestral alleles lost in the human OOA bottleneck or later migrations into the ancestors of Eurasian populations.

For the majority of RAs, the reintroduced allele is still segregating in African populations; however, a substantial fraction of RAs (EAS: 22%, EUR: 30%, and SAS: 28%) are present in modern human populations exclusively on haplotypes of Neanderthal ancestry (i.e., these alleles are no longer present in African populations). This suggests that the derived allele likely became fixed at these positions in AMH populations before the reintroduction of the ancestral allele via Neanderthal admixture. For those RAs where the corresponding allele is still present in Africans, they are segregating at significantly higher frequencies in Eurasians than those RAs no longer observed in Africans (**Figure** S6). This suggests heterogeneity in the selective pressures on RAs.

Next, we examined the distribution of RAs across introgressed haplotypes. RAs are pervasive; 84.4% (EAS), 81.8% (EUR), and 81.7% (SAS) of introgressed haplotypes contain RAs. The average number of RAs per introgressed haplotype is ∼17. (**Figure** S7A). Of the haplotypes containing RAs, 21.3% (EAS), 11.8% (EUR), and 15.2% (SAS) contain more RAs than NDAs (**Figure** S7B). RAs also have greater variability in their distribution across haplotypes, and appear more clustered within haplotypes than NDAs (**Figure** S7C). Thus, RAs are present on most introgressed haplotypes and, in some cases, constitute the majority of introgressed variants in these regions.

### RA-containing introgressed haplotypes are associated with anthropometric human traits and disease risk

To update knowledge of human phenotypes influenced by Neanderthal introgression, we intersected all RAs and NDAs from each of the three Eurasian populations with significant associations (*P* < 10^−8^; Methods) reported in the GWAS Catalog as of January 24, 2019 (*33*). Sixty-eight percent of NDAs with at least one significant GWAS association are in perfect LD with at least one RA (**File** S2). The consequence of this is that over 70% of the phenotype associations with NDAs have an equally strong association with at least one RA. Thus, while previous studies have used GWAS to link variants on introgressed haplotypes with phenotypes (*5, 6, 9*), many associations could be mediated by RAs.

The high LD between RAs and NDAs prevents the identification of the RAs, the associated NDAs, or other variants as causal. However, in Europeans, we found that nearly as many RAs (n = 1049) as NDAs (n = 1349) are significantly associated with at least one trait. Overall, Eurasian RAs tagged 2197 unique, significant associations while NDAs tagged 2547 (**File** S2). Many of the phenotypes tagged by RAs are morphometric (e.g., cranial base width, BMI, and height), and several others relate to more general aspects of outward appearance (e.g., chin dimples, male-pattern baldness, and skin pigmentation). Introgressed RAs are also associated with many pathologies, including cancers (breast, esophageal, lung, prostate), Alzheimer’s disease, and neurological conditions like neuroticism and bipolar disorder (**File** S2).

Several of the RAs that are no longer present within sub-Saharan African populations also have associations with traits. These RAs are particularly interesting, because they likely represent loci at which derived alleles became fixed in modern human populations after the split from ancestors of Neanderthals. For example, an RA (rs11564258) near *MUC19*, a gel-forming mucin expressed in epithelial tissues with a potential role in interaction with microbial communities, is strongly associated with both Crohn’s disease and inflammatory bowel disease (*34, 35*). This locus has been identified in scans for potential adaptive introgression (*18*). We also find associations with facial morphology, body mass index, sleep phenotypes, and metabolite levels in smokers (*36*–*40*). While none of these findings suggest that RAs are more likely than NDAs to be “causal”, they significantly expand the number of candidate variants in introgressed regions.

### Introgressed haplotypes containing eQTL have higher RA fraction than non-eQTL introgressed haplotypes

We next evaluated the prevalence of RAs among eQTL in 48 tissues profiled in v7 of the Genotype-Tissue Expression (GTEx) project (*41*). Introgressed eQTL are found in all GTEx tissues, with 18% of EUR RAs (16,318) and 16% of EUR NDAs (31,822) being eQTLs in at least one tissue. While each RA is associated with at least one NDA, the number of RAs and NDAs on an introgressed haplotype is not perfectly correlated (Pearson r^2^ = 0.46; Figure S7A). The 1585 introgressed haplotypes containing at least one introgressed eQTL have a significantly higher fraction of RAs than the 4237 haplotypes having no eQTL (median of 0.20 vs. 0.24, P = 3×10^−13^, Mann-Whitney *U* Test; **Figure** 3A). This result also holds when stratifying introgressed haplotypes with eQTL by their tissue of activity (**Figure** S14).

**Figure 3.**
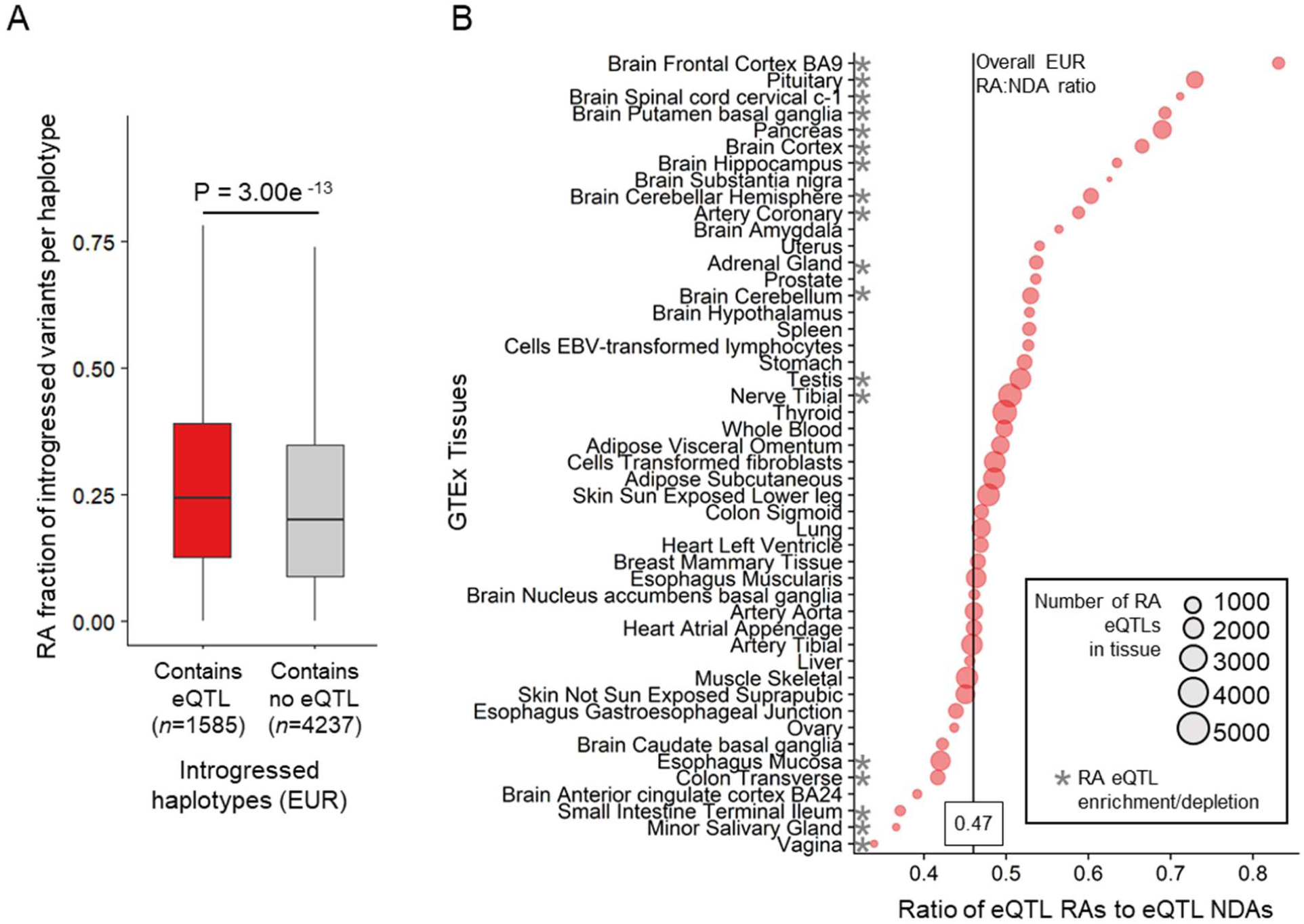
Reintroduced alleles are common among introgressed eQTLs. (A) The fraction of RAs among introgressed alleles on Neanderthal haplotypes in Europeans (EUR) that either contain or are lack GTEx eQTL. Introgressed haplotypes with eQTL have significantly higher RA fraction (median of 0.24 vs. 0.20, *P* = 3.00e–13, Mann-Whitney *U* test). (B) The RA:NDA ratio among eQTLs in each of 48 GTEx v7 tissues tissue. Bubbles are scaled by the number of RA eQTLs in each tissue. Compared to the genome wide average (RA:NDA ratio = 0.47; indicated by vertical black line), 13 tissues show more than the expected number of RA eQTL and 5 tissues show fewer than the expected number (*P* < 0.01, hypergeometric test after Bonferroni correction).

Among introgressed variants that are also eQTL, the ratios of RAs to NDAs varied across tissues, with 13 tissues having a higher RA:NDA ratio than expected from the RA:NDA ratio in the genome as a whole (**Figure** 3B). Brain tissues are 7 of the 13 tissues enriched for RA eQTLs, having RA:NDA ratios of between 0.53–0.83 compared to the overall observed ratio of 0.47 (P < 0.01, hypergeometric test with Bonferroni correction). Introgressed haplotypes have been previously shown to modulate gene regulation, especially in the brain (*24, 42*), and the higher-than-expected presence of RAs in more than half of these tissues could suggest shared regulatory architectures. RA eQTLs also appear more abundant in the pituitary gland, pancreas, adrenal gland, testes, and tibial nerve. RA eQTL are less abundant than expected in the introgressed eQTL from mucosal tissues and salivary gland. In summary, introgressed haplotypes containing eQTL contain a higher fraction of RAs, and this set of RA eQTLs is not evenly distributed among tissues.

### Some RAs have conserved gene regulatory associations in European and African populations

Many introgressed haplotypes influence gene regulation, and the majority of them contain RAs (*42, 43*). Given the high LD between RAs and NDAs, it is challenging to determine from genetic association data alone whether a particular RA or NDA is functional. Indeed, we find that RAs and NDAs are similarly likely to overlap known regulatory motifs (**Figure** S8). Thus, to search for RAs likely to have regulatory functions independent of associated NDAs, we analyzed cross-population eQTL data from lymphoblastoid cell lines (LCLs) from European (EUR) and sub-Saharan African Yoruba (YRI) individuals (*44*). We sought eQTL alleles that are RAs in Europeans and are present in Yoruba in non-Neanderthal introgressed regions (**Figure** 4A; Methods). Thus, if an allele that was reintroduced into Eurasians has similar effects on gene expression in both populations, it suggests that that the RA (rather than associated NDAs) influences expression, and that introgression reintroduced ancestral regulatory function.

**Figure 4.**
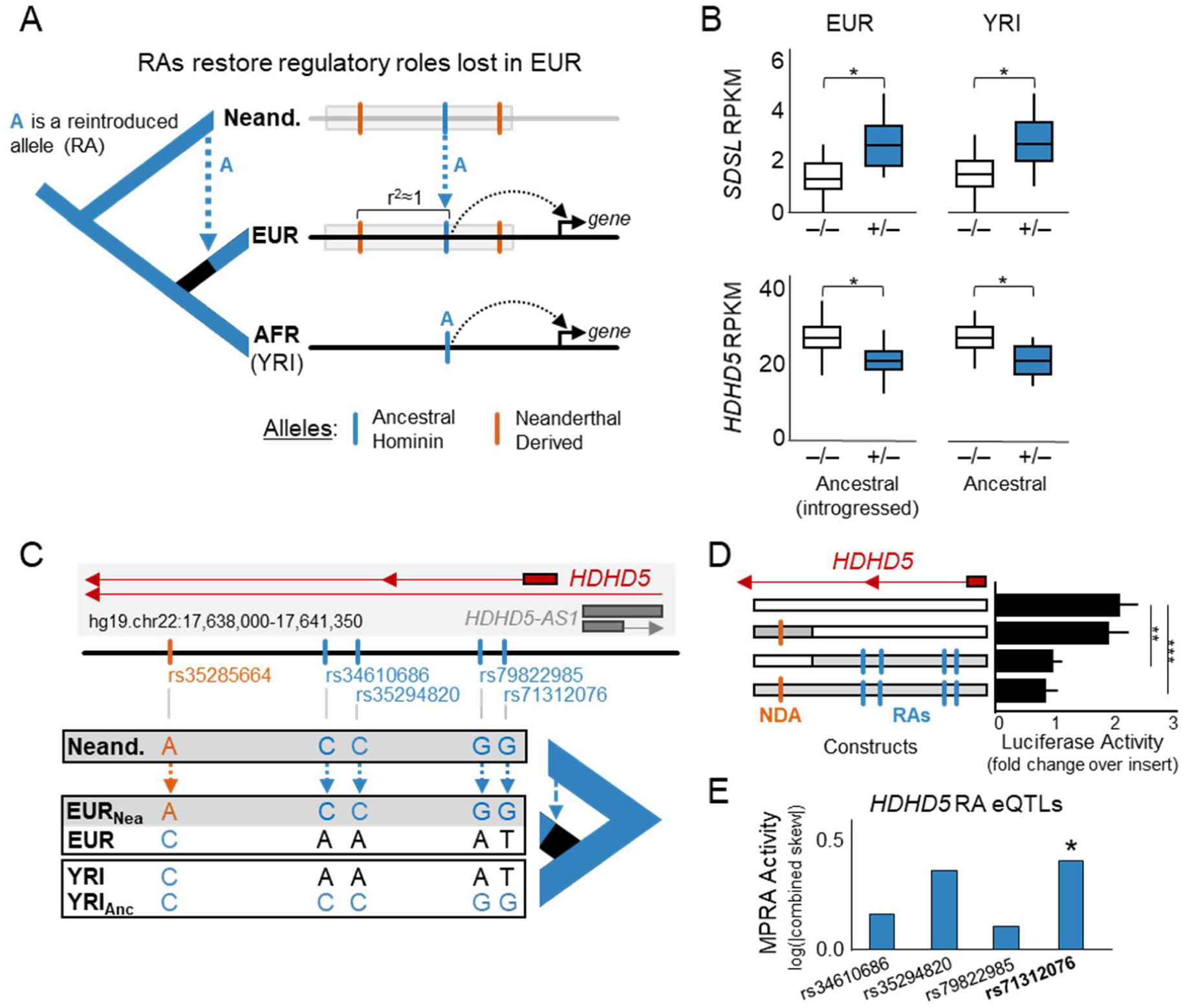
Reintroduced alleles restore regulatory functions lost in Eurasians. (A) Conceptual model of restored regulatory function resulting from Neanderthal admixture. Here, allele A is a *cis*-acting regulatory variant that is exclusively found on introgressed haplotypes (gray) in modern Europeans (EUR). Allele A is also present in sub-Saharan Yoruba individuals (YRI) lacking Neanderthal ancestry. It displays similar *cis*-regulatory activity in both populations. This pattern suggests that allele A is an RA in Europeans and that it influences gene regulation independent of the associated NDAs. (B) Two examples of genes (*SDSL* and *HDHD5*) with consistent expression differences (measured in RPKM) associated with RA eQTLs in EUR and the corresponding allele in YRI LCLs. The RAs are present only on introgressed haplotypes in EUR, and the NDAs associated with the RAs are not present in YRI. This suggests that these RAs restore lost gene regulatory functions in Europeans. (C) Schematic of the *HDHD5* locus highlighting the locations of one NDA (orange) and four RA eQTLs (blue) in the introgressed haplotype and the different combinations of these alleles present in EUR, YRI, and Neanderthals. (D) Luciferase activity driven by constructs carrying different combinations of alleles present in the *HDHD5* locus. We assayed four constructs containing: 1) no introgressed alleles, 2) only the NDA, 3) only the RAs, and 4) all introgressed variants. Results are summarized over three replicates. As expected from the eQTL data, constructs lacking RAs drive significantly stronger expression (∼2x baseline) than constructs containing RAs (∼1x baseline; two-tailed t-test, P < 0.01 (**) and P < 0.001 (***)). The regulatory effect of the RAs is independent of the presence the NDA found in introgressed EUR haplotypes. (E) Regulatory activity in a massively parallel reporter assay (MPRA) for the four *HDHD5* RA eQTLs reveals that rs71312076 has significant (P < 0.007) regulatory effects when placed in the non-introgressed European background sequence.

In the LCL eQTL data, 2,564 RAs were significant eQTL in EUR, and only 180 were significant eQTL in YRI. This difference is largely due to the much lower power in YRI (sample size of 89 vs. 379) which, in combination with having cross-population data from only one cellular context, makes it challenging to estimate the full extent to which RAs contribute regulatory function. Nevertheless, of the 180 YRI eQTL corresponding to EUR RAs, 42 displayed significant eQTL effects in both populations. These RA eQTLs influence the expression of nine genes (**Table** S3). The expression differences observed for the RAs in EUR have the same direction of effect and similar magnitude as those observed for the corresponding allele in YRI. For example, two genes, *SDSL* and *HDHD5*, each have four cross-population RA eQTLs that have similar effects on gene expression in both EUR and YRI (**Figure** 4B). Thus, despite the limitations of the cross-population eQTL data, these results suggest that some RAs influence gene regulation in Eurasian individuals.

### RAs can influence expression independent of NDAs

To determine whether RAs directly influence expression in EUR individuals independently of linked NDAs, we functionally dissected the regulatory activity of four cross-population RA eQTL. These alleles associate with the expression of *HDHD5* (also known as *CECR5*), a hydrolase domain containing protein that is expressed in diverse tissues. It is located in a region of chromosome 22 associated with Cat Eye Syndrome (CES), a rare disease associated with chromosomal abnormalities in 22q11 with highly variable clinical presentation that often includes multiple malformations affecting the eyes, ears, anus, heart, and kidneys (*45*). The *HDHD5* locus contains a 2 kb region, which in introgressed Europeans carries an NDA that is in perfect LD with four RAs that are cross-population eQTLs for *HDHD5* (**Figure** 4C).

We performed luciferase reporter assays in LCLs using four different combinations of the NDA and RAs within this 2 kb region (**Figure** 4D, **Table** S5). In each assay, we compared the activity of each combination of alleles to the activity driven by a vector with a minimal promoter but no insert. The luciferase activity driven by a reporter construct with the European version of this sequence without introgression (EUR-EUR) drove significant expression above baseline (∼2.0x vector with no insert, *P* < 0.01, t-test). We compared this activity to constructs synthesized to carry the RAs with the associated NDA (NDA-RA), the RAs without the NDA (EUR-RA), and the NDA without the RAs (NDA-EUR). Both RA-containing sequences had significantly lower luciferase activity, and there was no significant difference in the activity of the NDA-RA and the EUR-RA sequences (**Figure** 4D). Thus, as predicted by the cross-population eQTL data, the RAs are associated with expression levels independently of the NDA, and the RA-containing sequences have lower activity than sequences without the RAs.

To ascertain whether the conservation of activity patterns we demonstrated at the *HDHD5* locus could be attributed more specifically to any of the four RAs, we analyzed previously collected MPRA data from LCLs (*46*). Only one of the four cross-population RA eQTL (rs71312076) showed significant regulatory effects (RA:EUR allelic skew=2.122, *P*=6.6e-3, FDR=0.034) compared to the non-reintroduced allele (**Figure** 4E). These effects were observed on the non-introgressed European reference background, further demonstrating the ability of this RA locus to influence regulation independent of NDAs.

Together, these results provide three orthogonal lines of evidence (cross-population eQTL, luciferase reporter, and MPRA) implicating RAs in the reintroduction of regulatory effects in the *HDHD5* locus. Importantly, both our luciferase assays and the MPRA data show that the functional contribution of RAs within a European genomic context is not dependent on the introgressed haplotype in which it occurs. Therefore, these data, along with the eQTL status of this region in YRI, demonstrate that Neanderthal introgression restored an allele lost in Eurasians that influences gene regulation.

## DISCUSSION

Here we demonstrate that hundreds of thousands of ancient alleles are present in modern Eurasians due exclusively to archaic admixture between Neanderthals and AMHs (**Figure** 1A). We first show that like NDAs, RAs are as associated with many traits and are enriched on haplotypes with regulatory effects in some tissues. We further show that RAs can have gene regulatory functions that are not dependent upon linked NDAs. While the interpretation of the phenotypic effects of Neanderthal introgression on AMHs has generally focused on NDAs, our results argue that RAs have the potential to independently affect gene regulation and therefore must also be considered in analyses of archaic admixture.

The evolutionary histories of RAs are likely diverse. While most RAs were probably lost in the Eurasian OOA bottleneck, Eurasian subpopulations subsequently experienced distinct demographic events that could have led to population-specific RAs. For example, East Asians are estimated to have had both substantially smaller ancestral effective population size than Europeans, as well as a greater frequency of archaic introgression events (*32*). These factors would increase our power to detect RAs within East Asians, because ancient hominin alleles would have had more opportunities to both be lost and to occur exclusively within introgressed regions. Conversely, in southern Europe, where more recent gene flow out of Africa has occurred, the power to detect RAs will be decreased due to ancient alleles being re-introduced outside of introgressed haplotypes (*47*). We expect that more sophisticated simulations and probabilistic modeling could enable the identification of additional RAs.

The regulatory and phenotypic effects of RAs are difficult to disentangle from those of NDAs, due to their high LD. Our detailed *in vitro* analyses of different combinations of alleles at the *HDHD5* locus (**Figure** 4D) provides a roadmap for experimental characterization of other RAs. Analysis of known regulatory elements suggests that at least 10% (19,882) of RAs overlap gene regulatory elements (**Figure** S8). Therefore, we anticipate that as MPRAs, eQTL analyses, and GWAS are performed in more diverse populations and tissues, more functional RAs will be identified. In the future, it will also be informative to compare the functional effects of RAs with other alleles restored to Eurasian populations more recently by direct migration from Africa (*22, 48*).

Given our *in vitro* demonstration that RAs can restore ancestral gene regulatory functions lost in Eurasian populations, the enrichment we observe for RAs relative to NDAs in some GTEx tissues—the brain in particular—is provocative (**Figure** 3B). These observations are consistent with previous results regarding the gene regulatory effects of introgressed alleles. Brain tissues have enrichment for Neanderthal eQTL (*24*), and there is significant allele-specific down regulation of haplotypes carrying Neanderthal alleles in the brain and testes (*42*). Our results add to this picture, suggesting that RAs may contribute to the regulatory architectures of some tissues.

Several non-exclusive evolutionary scenarios may explain these observations. First, the depletion of NDAs relative to RAs on introgressed haplotypes with gene regulatory functions could be a result of previously demonstrated selection against NDAs in some tissues (*42*). This selection would deplete tissue-specific regulatory regions of NDA-rich introgressed haplotypes; indeed, the two tissues with known allele-specific down regulation of Neanderthal alleles, brain and testes, are among those enriched for RAs compared to NDAs. Second, the patterns we see could result from positive or balancing selection acting to retain beneficial RAs. Under this scenario, archaic admixture restored alleles with beneficial regulatory functions that were lost to Eurasian populations, and these RAs contributed to the maintenance of some introgressed haplotypes. The third possibility is that both RAs and NDAs on introgressed haplotypes are functional and influence selective pressures on the haplotypes. In this case, the presence of RAs could counterbalance mildly deleterious effects of associated NDAs, and thus buffer some introgressed haplotypes from purifying selection. Importantly, these explanations are not mutually exclusive, and the reality is likely some combination of all of them.

Overall, we anticipate that the regulatory effects of RAs and NDAs differ between tissues based on the genetic diversity and strength of constraint on their regulatory landscapes. Supporting this, nervous system tissues (including the brain) and the testes have extreme levels of selection on gene expression (high and low, respectively) (*49*). Given the range of RA eQTL enrichments across GTEx tissues, including tissues without evidence of selection against Neanderthal alleles, we propose that the presence of RAs and NDAs is the result of a mixture of selective pressures acting within the regulatory constraints of each tissue. Therefore, RAs likely play a functional role across diverse tissues and thus may contribute to the persistence of introgressed haplotypes.

Contrasting the functional effects of RAs and NDAs will be especially useful in understanding how evolution acted on introgressed haplotypes. Although RAs and NDAs co-segregate in Eurasians, they are the products of two distinct evolutionary histories. Previous work has implicated the small effective population size of Neanderthal populations as a key factor in their transmission of weakly deleterious NDAs into AMHs via introgression (*19, 20, 50*). In contrast, RAs are more evolutionarily conserved than the NDAs, arose in a genomic background ancestral to AMHs, and were maintained in a relatively larger ancestral hominin population in which selection could act more efficiently. Accordingly, we expect RAs and NDAs to have different distributions of functional effects in introgressed human populations. While estimating effects of individual mutations is challenging, variant effect prediction algorithms (CADD and PolyPhen2) indicate that NDAs are generally more damaging than RAs (**Figure** S9, **Figures** S11-S13), consistent with a model of less efficient selection on NDAs than RAs over their histories. Consequently, coding or regulatory functions played by RAs should be more benign overall than those of NDAs, perhaps even beneficial. Thus, within the context of some introgressed haplotypes, the ancient RAs may have conferred a degree of “compatibility” that protected the linked NDAs from negative selection.

Further analysis of RAs will also be relevant to studies of the genetics of ancient hominin populations. For example, tens of thousands of RAs that are present in Eurasians are not present in African populations. These ancient variants could both inform ongoing debates over differences in efficiency of natural selection between Africa and Eurasia (*51*–*54*), as well as provide a window into ancient genetic variation that was present in Africa over a half million years ago.

## CONCLUSIONS

Here we show that Neanderthal introgression reintroduced alleles lost in to the ancestors of Eurasian populations and that many of these RAs have the potential to be functional. This illustrates the importance of accounting for shared ancestral variation among hominin populations and shows that hybridization events between populations have the potential to modulate the effects of bottlenecks on allelic diversity. Our findings open several avenues for future work on quantifying the evolutionary and functional dynamics of archaic introgression. Previous analyses of introgression have focused on alleles derived within the Neanderthal lineage; additional work is needed to account for the potential effects of RAs and their influence on the maintenance of Neanderthal ancestry. In short, reintroduced alleles must also be considered in analyses of Neanderthal introgression, at both the haplotype and genome scale.

## ACKNOWLEDGMENTS

We thank Ben Haller, Phillip Messer, and Kelley Harris for advice on evolutionary simulations. We thank Ryan Tewhey for discussions of MPRA results. This work was supported by the National Institutes of Health: T32EY021453 to CNS; T32GM080178 to DS; K22CA184308 to EH; and R01GM115836 and R35GM127087 to JAC. This work was conducted in part using the resources of the Advanced Computing Center for Research and Education at Vanderbilt University, Nashville, TN.

## AUTHOR CONTRIBUTIONS

DCR, CNS, EM and JAC conceived and conducted the computational analyses. DS and EH performed the luciferase assays. DCR and JAC wrote the manuscript with input from all authors.

## METHODS

### Sequence data

Genomic variants were taken from 1000 Genomes Phase 3v5a data (*1*). Introgressed Neanderthal tag SNPs were downloaded from: http://akeylab.princeton.edu/downloads.html (*12*). All analyses were conducted using GRCh37/hg19 genomic coordinates.

### RA candidate identification and classification from 1000 Genomes data

To generate a set of candidate RAs, we gathered Neanderthal tag SNPs identified in each of the three 1000 Genomes Eurasian super-populations (EUR, EAS, SAS). These tag SNPs represent variants that are rarely or never observed in African populations, yet that are present in Neanderthal introgressed haplotypes in Eurasians. We then calculated LD using vcftools (*55*) for all variants in +/–500 kb windows around each variant across individuals from these super-populations in Phase 3 of the 1000 Genomes project. We extracted all variants that were in perfect LD (r^2^=1) with any Neanderthal tag SNP in any of EUR, EAS, or SAS populations.

For each candidate RA (i.e., a variant in perfect LD with a Neanderthal tag SNP that was not itself a Neanderthal tag SNP) we: 1) extracted the ancestral allele call from 1000 Genomes, 2) ascertained whether the designated REF or the ALT allele was the introgressed variant (i.e., in LD with the Neanderthal tag SNP), 3) calculated the introgressed allele frequency, 4) calculated the allele frequency for the same allele in sub-Saharan African 1000 Genomes populations, and 5) extracted the Altai Neanderthal genotype. We then assigned RA status based on this information by following to the steps laid out in **Figure** S3. Specifically, for each RA candidate, if the introgressed variant matched the high-confidence, ancestral state, it was classified as a reintroduced ancestral allele (RAA). Candidate RAs that did not match the ancestral allele (or that did not have a high confidence ancestral allele call) were evaluated for presence in both the Altai Neanderthal and in sub-Saharan Africans (average allele frequency > 1% in ESN, GWD, LWK, MSL, and YRI). If the candidate variant was only present in sub-Saharan African at a frequency > 1%, it was classified as a reintroduced hominin alleles (RHA) since its origin likely predated the Neanderthal split, but its ancestral status is not assigned. Importantly, the criteria that an RHA also be present at a minimum allele frequency of 1% frequency over all 5 sub-Saharan populations insulates our results from the trace levels of Neanderthal ancestry from Eurasian backflow (0.18% Neanderthal ancestry in YRI) (*29*). If the candidate variant was only present in the Altai Neanderthal and in introgressed Eurasian haplotypes, it was classified as an NDA. For nearly all analyses presented here, RAAs and RHAs are combined into a single RA class. The results of this classification are summarized in **Figure** 2 and supplied in full in **File** S1. The pipeline and filtering steps are summarized in **Figure** S3.

We considered all Neanderthal haplotypes and tag SNPs identified by Vernot et al. 2016. Only 10% of identified RAs were found in haplotypes shorter than 10 kb or with fewer than 10 NDAs. Removal of these NDAs and RAs from our analyses did not substantially change any of our results, thus we present results with all RAs for completeness.

Approximately 90% of RAs are within the boundaries of previously characterized introgressed haplotypes; however, over half of the haplotypes in each population have at least one associated RA beyond their previous bounds. In total, extending all introgressed haplotypes to accommodate all associated RAs increases introgression estimates by 40.0, 42.6, and 51.9 megabases (Mb) in the EUR, EAS, and SAS populations, respectively. This represents an increase of ∼1.5% in the amount of introgressed sequence present in each Eurasian population.

As described above, when no confident ancestral allele call was available, we used the presence of an allele in modern Africans to infer that it was present in the ancient hominin population. In such cases, if the allele was present in modern Eurasians on a Neanderthal haplotype, it was inferred to be an RHA. However, without a confident ancestral state for these alleles, RHAs may be susceptible to false positives due to independent, convergent mutations on the AMH African and Neanderthal lineages. This is of particular concern at CpG sites due to their significantly higher mutation rates compared to other dinucleotides. Therefore, to estimate how many inferred RHAs could be the result of independent C→T transitions along the Neanderthal and AMH lineages, we identified all RHAs that were either “A” or “T” with a 5’ “C” or 3’ “G” respectively. We focused on RHAs, because RAAs match ancestral allele calls supported by cross-species alignments making convergent mutation unlikely. Only 7% of RHAs were potentially subject to this bias (**Table** S7). Furthermore, we carried out simulations with an estimate of the CpG mutation rate (7.0 × 10^−7^ mutations per site per generation (*56*)) and estimated that 3% of all CpGs would be expected to display convergent mutations between Neanderthals and AMH Eurasians. Thus, confounding due to convergent mutations, even at CpGs, is likely to be rare (**Table** S7). (See “Estimating confounding with simulations” section for more details on the simulations.)

### Spatial characterization of RAs and NDAs along introgressed haplotypes

The locations and distributions of RAs within introgressed haplotypes are less correlated with haplotype length and more clustered than the distribution of NDAs. The number of NDAs per haplotype is strongly positively correlated with the length of the haplotype (r^2^ = 0.85; **Figure** S7), but the RA number per haplotype is more variable (r^2^ = 0.56). Therefore, while the overall RA:NDA ratio is ∼1:2 over all haplotypes (**Figure** 2), this ratio varies across introgressed haplotypes.

To evaluate whether RAs are more clustered on introgressed haplotypes than NDAs, we summarized the distribution of both NDAs and RAs across all RA-containing haplotypes. We first divided each RA-containing haplotype into 100 equal-size bins and counted the number of RAs in each bin. For each haplotype, the bins were then ranked from high to low in terms of RA count, and the RA contents of each corresponding percentile bin were summed over all the haplotypes. This percentile sum was then divided by the total number of all RAs present over all the haplotypes to obtain per-bin densities. By calculating per-bin densities only at the end, we mitigate the potentially confounding effect of some haplotypes containing fewer variants than others. The result is a summary of the total fraction of RAs found within increasing density percentiles across all haplotypes. We then did the same for NDAs (**Figure** S7) Overall, a larger fraction of RAs is found in the densest bins compared to NDAs. For example, in EUR, 55% of RAs are in the four densest bins, while only 26% of NDAs are in the four densest bins. These results held across each population and were maintained when down sampling to a set of haplotypes with matched NDA and RA counts. Thus, when RAs are present, they often occur in more discrete clusters along introgressed haplotypes than do NDAs. However, we note that the incomplete ascertainment of RAs and the LD thresholds used to link NDAs may contribute to these patterns.

### Computational variant effect estimation

To assess the potential functional impact of RAs, we analyzed precomputed Combined Annotation-Dependent Depletion (CADD) v1.3 scores (https://cadd.gs.washington.edu/download) for all RA and NDA variants. CADD scores are available in two forms: raw and scaled. Raw CADD scores for variants are the output of a support vector machine trained to distinguish variants observed in 1000 Genomes (likely benign) from non-observed variants (likely deleterious) based on diverse annotations. To enable comparisons across sites, the raw scores for all possible mutations to the hg19 genome were ranked and PHRED-scaled (–10 * log_10_(*Rank*)) (*57*). Therefore, the scaled scores communicate how deleterious the effect of a given variant is with respect to the effects seen in all other possible variants (e.g., a scaled CADD of 20 means that that a variant is within the top 1% of variants as ranked by their predicted deleteriousness). Thus, we focused on the PHRED-scaled scores. In particular, we highlighted in **Figure** S9, **Figure** S11, and **Figure** S12, scaled CADD scores at the upper range (e.g., above 10 or 15) that are suggestive of deleteriousness. We also compared functional annotation classes downloaded for RAs and NDAs from RegulomeDB v1.1 (**Figure** S8; http://www.regulomedb.org/) and PolyPhen2 (**Figure** S13; http://genetics.bwh.harvard.edu/pph2/).

### Evaluation of introgressed variants in GWAS Catalog

To evaluate the prevalence of RAs among significant GWAS associations, we intersected all RAs and NDAs with variants reported in GWAS Catalog (as of January 24, 2019). To account for other variants tagged by the variant reported in the GWAS Catalog, we expanded the target set of GWAS variants to include variants in perfect LD (1000 Genomes r^2^=1) in each of the three Eurasian populations. We then intersected sets of introgressed variants from each population with LD-expanded GWAS hits for that population. The results of these expansions and intersections are presented in **File** S2.

### GTEx eQTL enrichment analysis

Expression quantitative trait loci (eQTL) data from GTEx v7 were downloaded from the GTEX portal (https://www.gtexportal.org/home/datasets) and all significant gene-eQTL pairs were extracted for each tissue. We then identified all RAs and NDAs with eQTL status. Because, the GTEx cohort is >85% European ancestry, we only considered European RAs and NDAs in this analysis.

First, we evaluated whether introgressed haplotypes with eQTL activity have a different RA fraction than introgressed haplotypes without eQTL (**Figure** 3A). This observation held when eQTL were stratified by tissue (**Figure** S14), suggesting that at the haplotype level, RAs appear at greater frequency within regions associated with gene regulation.

Next, within each tissue, we considered only those significant introgressed (i.e. RA and NDA) eQTL and the RA:NDA ratio for each tissue in **Figure** 3B. To test whether there was an overrepresentation of RAs among introgressed eQTLs, we performed a hypergeometric test on each tissue’s set of introgressed eQTLs with respect to the background of all variants evaluated for eQTL status within GTEx. We applied a Bonferroni correction to account for the testing of the 48 tissues (0.01/48=0.0002).

### Shared RA eQTLs between Europeans and Africans

To identify RAs with similar regulatory associations between populations with and without Neanderthal ancestry, we analyzed data from a previous study that identified eQTL across LCLs derived from 495 individuals (*44*). The LCLs were of either European (EUR; 373 lines) or African (YRI; 89 lines) ancestry; given the smaller YRI sample size, there was much lower power to detect eQTL in the African samples. We downloaded all significant exon-level expression eQTLs from the study (https://www.ebi.ac.uk/arrayexpress/files/E-GEUV-1/analysis_results/). They found 704,157 unique eQTL in EUR and 75,742 in YRI, and of these, 52,869 are shared. Of the shared loci, 42 are RAs, and these RAs associate with expression levels for nine genes (**Table** S3). For each of the 42 variants, we also confirmed that they were not in LD in YRI with any previously characterized NDA.

### MPRA analysis of RAs

A recent MPRA study evaluated the regulatory impact of 32,373 variants in 3,642 known eQTL and regions identified via GWAS (*46*). For each variant, the MPRA quantified the expression of a reporter driven by both the reference and alternate alleles (plus 150 bp of reference genomic context) in LCLs. Expression modulating variants were identified by quantifying the “allelic skew” between the expression driven by the reference and alternate allele. This enabled the identification of hundreds of variants likely to cause the observed associations between these loci and expression levels/phenotypes. We intersected European NDAs and RAs in introgressed haplotypes with the variants with significant combined skew (FDR < 0.1). In total, 11 introgressed variants were tested (6 NDAs and 5 RAs; **Table** S4). This included all cross-population RA eQTLs in the introgressed haplotype that is associated with *HDHD5* expression.

### Experimental validation of RA regulatory function via luciferase assays

To further demonstrate that the cross-population RA eQTLs associated with *HDHD5* expression function independently of the NDA in perfect LD, we evaluated the effects of four different sequences on luciferase expression in LCLs (**Figure** 4D).

Modified pGL4 luciferase constructs were generated via Gibson cloning (New England Biolabs) to contain an 1826 bp oligo corresponding to region of interest in *CECR5/HDHD5* with variants corresponding to a European reference (EUR-EUR), the introgressed NDA sequence (NDA-EUR), the RA sequences (EUR-RA), or both sets of introgressed variants (NDA-RA) (**Table** S5). Inserts were cloned into the pGL4.27 reporter vector (Promega) as two separate blocks, as b1-EUR or b1-NDA (first 576 bp at the 3’ end of blocks containing either NDA or EUR specific sequence) and b2-EUR or b2-RA (1273 bp at the 5’ end of blocks containing either RA or EUR specific sequence) (**Table** S5). b1-EUR, b1-NDA, and b2-RA sequences were generated by oligonucleotide synthesis (IDT). b2-EUR variants were generated via site-directed mutagenesis using primers with EUR specific alleles (**Table** S6) and amplified directly from the b2-RA oligo as five separate sub-regions. B2-EUR sub-regions were assembled into the pGL4.27 vector and sub-cloned into EUR-EUR and NDA-EUR pGL4 constructs. Inserts were amplified to include NheI and XhoI overhangs to allow for cloning into the pGL4 reporter plasmid. The sequences of full-length inserts were confirmed by Sanger sequencing (Genewiz).

GM11831 B-cells were cultured in RPMI with penicillin/streptomycin and 15% fetal bovine serum. 1×10^6^ GM11831 cells were transfected with 5 ug HDHD5-EUR-EUR-pGL4.27, HDHD5-NDA-EUR-pGL4.27, HDHD5-EUR-RA-pGL4.27, or HDHD-NDA-RA-pGL4.27 along with 500 ng pRL-CMV (Renilla reporter plasmid) via electroporation (Neon Transfection System, Invitrogen). Firefly and Renilla luciferase activity were analyzed using the Dual-Glo Luciferase Assay System (Promega) and Synergy HTX MicroPlate Reader (BioTek) 19 hours post electroporation. Firefly reporter expression was normalized to Renilla luciferase activity. Statistical significance was determined through a two tailed t-test comparing fold change of the normalized luciferase activity over an unmodified (no insert) pGL4.27 reporter control.

### Evolutionary simulation framework

We carried out evolutionary simulations to explore RA dynamics and estimate false positive rates under a range of scenarios. SLiM (v2.6) was used for all evolutionary simulations (*58*). We based our simulations on genomic and demographic models used in previous simulation studies of Neanderthal introgression and mutation load (*20*). In brief, the human genome was represented by a syntenic, locus-based model that reflects the gene structures in the hg19 reference genome. Nucleotide positions of exons were modeled individually while intergenic regions and chromosomal boundaries were modeled as single sites. We used a recombination rate of 1.0 × 10^−8^ crossovers per site per generation with probabilities in intergenic regions scaled by their respective sizes. Chromosome boundaries had a recombination rate of 0.5. To estimate false positives, we also considered each of three mutations rates: 7.0 × 10^−9^, 7.0 × 10^−8^, and 7.0 × 10^−7^ mutations per site per generation. The highest rate was included to simulate the high mutability at CpG dinucleotides, while the lowest is in keeping with genome-wide estimates for non-synonymous sites in humans. We simulated fitness effects (FE) of mutations based either on neutrality (FE=0) or purifying selection (FE drawn from gamma distribution with shape parameter 0.23 and mean selection coefficient −0.043) (*59*). We also considered Eurasian–Neanderthal admixture fractions of 0.02 and 0.04.

The general demographic model used is illustrated in **Figure** S1. Genetic diversity within the ancient human population (10,000 diploid individuals) was first established by allowing mutations to arise and evolve through a “burn in” period of 44,000 generations in the ancestral hominin population prior to subsequent migrations. To track allelic loss and reintroduction, we focused on segregating sites that were present in this simulated ancestral population immediately before the split between the human and Neanderthal lineages; we tracked all of these ancestral hominin alleles over the 18,000 subsequent generations that encompassed both the Neanderthal and Eurasian OOA bottlenecks. The ancestral Neanderthal population was subsampled to 1,000 individuals and both human and Neanderthal populations evolved separately for 16,000 generations (400,000-464,000 years assuming a generation time of 25–29 years).

The Eurasian OOA migration and Neanderthal admixture were then modeled as a simultaneous, discrete event that resulted in an admixed Eurasian population size of 1861 individuals (*20, 60*). The admixed Eurasian population was then allowed to evolve for 2000 generations before undergoing exponential growth leading to 20,310 modern Eurasians. One hundred replicates for both neutral or purifying selection models were run to evaluate properties of RAs and estimate rates of confounding mutations (**Figure** S2). To enable further analyses, snapshots of alleles present in each population were collected at four relevant timepoints for each simulation: t1) Neanderthal OOA, t2) immediately prior to the Eurasian migration, t3) immediately following admixture, and t4) modern human populations. Mutation origin was used to establish when (generation) and where (genome location and population) a variant arose and to trace its presence/absence through successive timepoints.

To quantify the frequency of RAs in simulated modern Eurasian populations we defined “ancestral hominin variants” as those alleles segregating in the simulated population immediately prior to the Neanderthal split ∼500,000 years ago (t2). We tracked segregating ancestral variants through the Neanderthal lineage and into the modern Eurasian population. We used SLiM’s mutation identifiers to track these ancestral variants through Neanderthals and into modern Eurasians over each replicate. We identified all the ancestral variants that passed into AMH exclusively through either 1) the Eurasian OOA migration or 2) archaic admixture with Neanderthals (RAs). We extracted allele counts and selection coefficients (in admixture models run with purifying selection) for these RA variants from the SLiM output. We then did the same for the simulated NDAs, the only other class of variants that entered the modern Eurasian populations exclusively through Neanderthal introgression. These data are summarized and contrasted in **Figure** S2 and **Figure** S9A.

### Estimating confounding factors with simulations

We explored several sources of possible confounding through simulation. First, we estimated the rate at which variants could be mis-assigned RA status as the result of independent, convergent origins in African and Neanderthal populations. To infer the frequency of such confounding events, all variants in simulated human and Neanderthal populations were compared immediately prior to admixture (t2) in each of the 100 replicates for each model. We chose 100 replicates due to the computational cost of each simulation and the fact that the variance in the output statistics stabilized well before reaching 100 replicates. Confounding variants were identified based upon a shared genomic location between existing variants in Africans and variants that arose within the Neanderthal lineage. These counts (false positives) were then contrasted with the number of non-Neanderthal derived mutations (true negatives), and found to be very rare (**Figure** S2). Moreover, because SLiM does not consider nucleotide state and allows for “stacked” mutations (i.e., mutations at the same locus), our estimates of false assignment of RA status in this model are conservative because we also considered nucleotide state in the real data. We also considered a mutation rate an order of magnitude higher than the genome-wide average (7.0 × 10^−7^ mutations per site per generation) to reflect the hypermutatbility of CpG dinucleotides (**Figure** S2B).

Second, it is also possible that non-Neanderthal alleles could have recombined on to introgressed haplotypes and subsequently been lost outside of the introgressed context. We reasoned that this scenario would be very unlikely, especially given our requirement of perfect LD between RAs and NDAs in modern Eurasian populations in the inference of RA status. To test this, we examined each of the simulated Eurasians (t4) and extracted all variants in perfect LD with an NDA in modern Eurasians. We then queried the simulation data from t2 to count how many of these candidate RA variants were not present on a Neanderthal haplotype. These variants in perfect LD with an NDA in modern Eurasians that were not present in Neanderthals (and that had not independently appeared within Eurasians) would be incorrectly inferred to be RAs by our approach. As expected, these events were very rare (1% of RAs or fewer) for each admixture fraction (**Table** S2). Furthermore, these are likely overestimates since in the real data, RAs most frequently appear within introgressed haplotypes, with linked NDAs present on both sides. This would suggest two recombination events, with all the confounding alleles then being subsequently lost on all non-introgressed haplotypes to maintain perfect LD. In the future, we anticipate that these simulations can be refined to confidently identify more RAs that retain lower LD with NDAs.

### Data analysis and visualization

Evolutionary simulations and primary data analysis were conducted on Vanderbilt’s computing cluster (ACCRE). Results were parsed and analyzed with custom python and bash scripts. Statistical tests were performed with R. Plots were generated in R, with most generated using ggplot2.

### Data availability

All results reported in this paper will be made available in supplementary material and/or on the project’s github repository (https://github.com/DaRinker/Neanderthal.reintroduction) subsequent to acceptance of the manuscript.

### Code availability

Full code used in this analysis will be available on the project’s github (https://github.com/DaRinker/Neanderthal.reintroduction) subsequent to acceptance of the manuscript.

## SUPPLEMENTARY FIGURES

**Figure S1.**
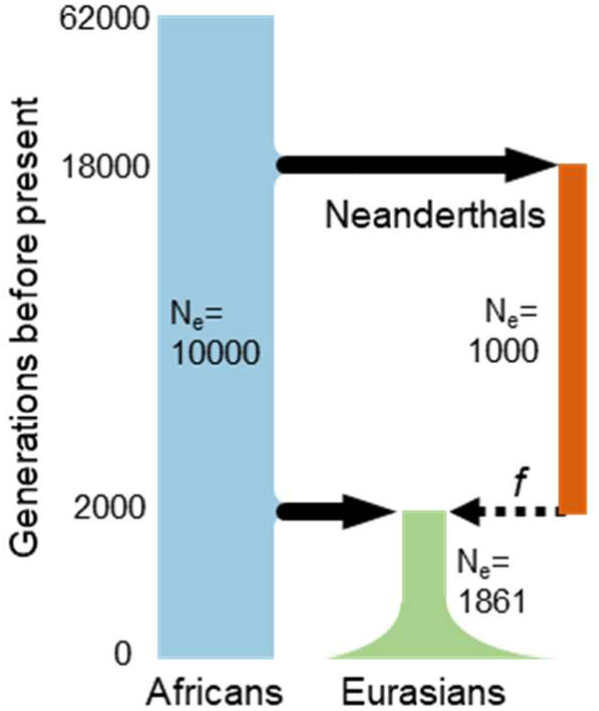
Demographic model used for evolutionary simulations. The demographic model used to simulate human–Neanderthal admixture and quantify the reintroduction of lost alleles. The model and effective population sizes (Ne) were based on previous simulations of Neanderthal admixture. We considered models in which mutations incurred a fitness cost (mildly purifying selection) or no fitness cost (strict neutrality). Two different admixture fractions (*f*=0.02 and *f*=0.04) and three mutation rates were used in the simulations (Methods).

**Figure S2.**
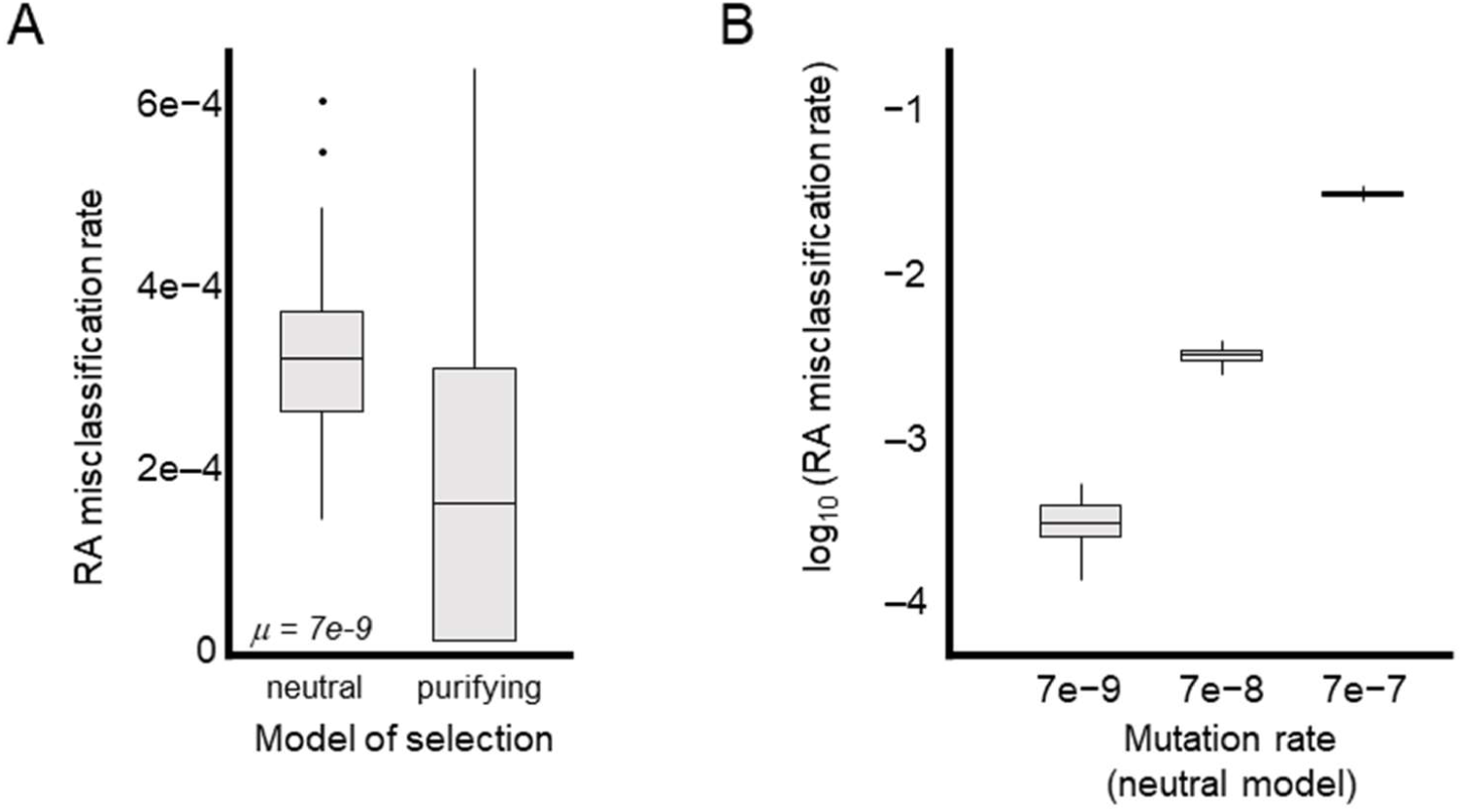
Simulations indicate that false positives in RA identification due to independent convergent mutations are rare. For each simulated population, we identified all NDAs that occurred in positions with ancestral hominin variation that was lost in the Eurasian OOA. (A) Boxplots summarize the frequencies of these potentially confounding NDAs among all sites that would be called as RAs at the time of admixture (c.f. **Figure** 1). The incidence of these confounding mutations is slightly higher under a purely neutral model (left) than under a model where new mutations can be deleterious (right). (B) Comparison of the effect of elevated mutation rates on the incidence of potentially confounding variants. Under a neutral model, the false positive rate scales with the mutation rate. The highest rate (μ= 7e-7) provides an estimate for CpG sites and results in a 3% false positive rate. Each boxplot represents 100 simulated populations.

**Figure S3.**
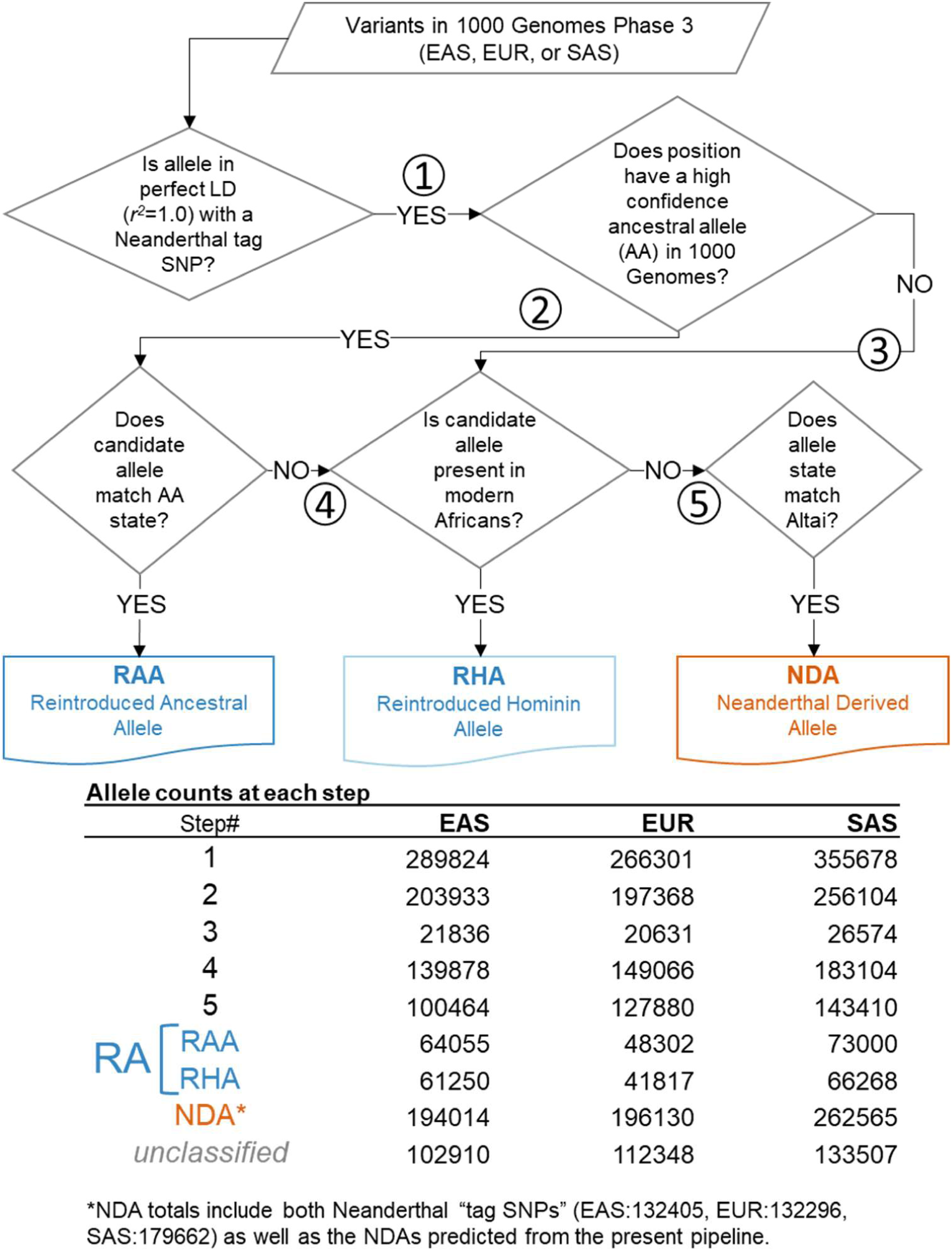
Introgressed allele class assignment decision tree and allele count summary. Decision tree by which 1000 Genomes variants in perfect LD with Neanderthal tag SNPs were classified as RAs and NDAs. The counts of variants making it to each of the numbered steps (1-5) is summarized in the lower table.

**Figure S4.**
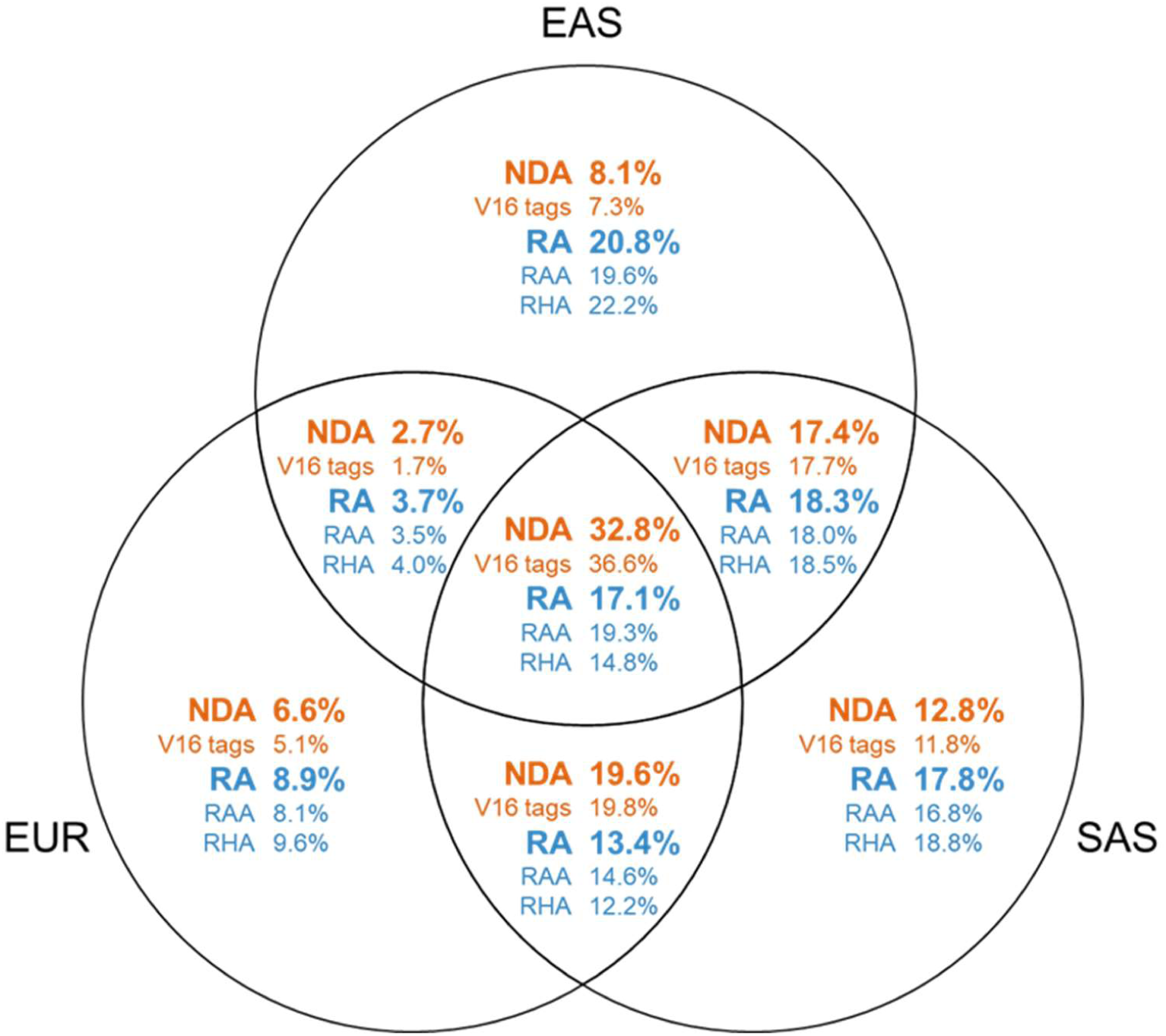
Introgressed allele sharing across three Eurasian populations. Venn diagram showing the fractions of each introgressed variant class that are shared between populations.

**Figure S5.**
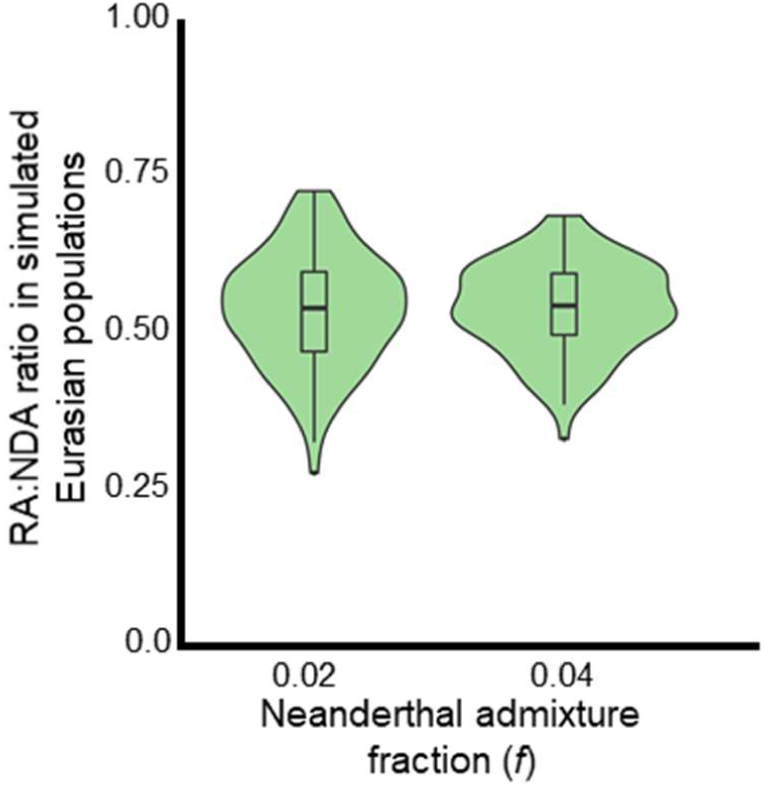
Simulations indicate that reintroduction of alleles lost in the OOA bottleneck by Neanderthal introgression was common. The ratios of RAs to NDAs over 100 simulated Eurasian populations. The simulations predict approximately one RA for every two NDAs, and these estimates are robust to changes in the simulated Neanderthal admixture fraction. Misclassification of non-RAs as RAs due to independent, convergent mutations is extremely rare (**Figure** S2) and the overall false discovery rate for LD-based RA identification is below ∼1% (**Table** S2). While these forward time simulations only approximate the demographic histories of these populations, the observed RA-to-NDAs ratio are qualitatively consistent with the simulations (**Figure** 2).

**Figure S6.**
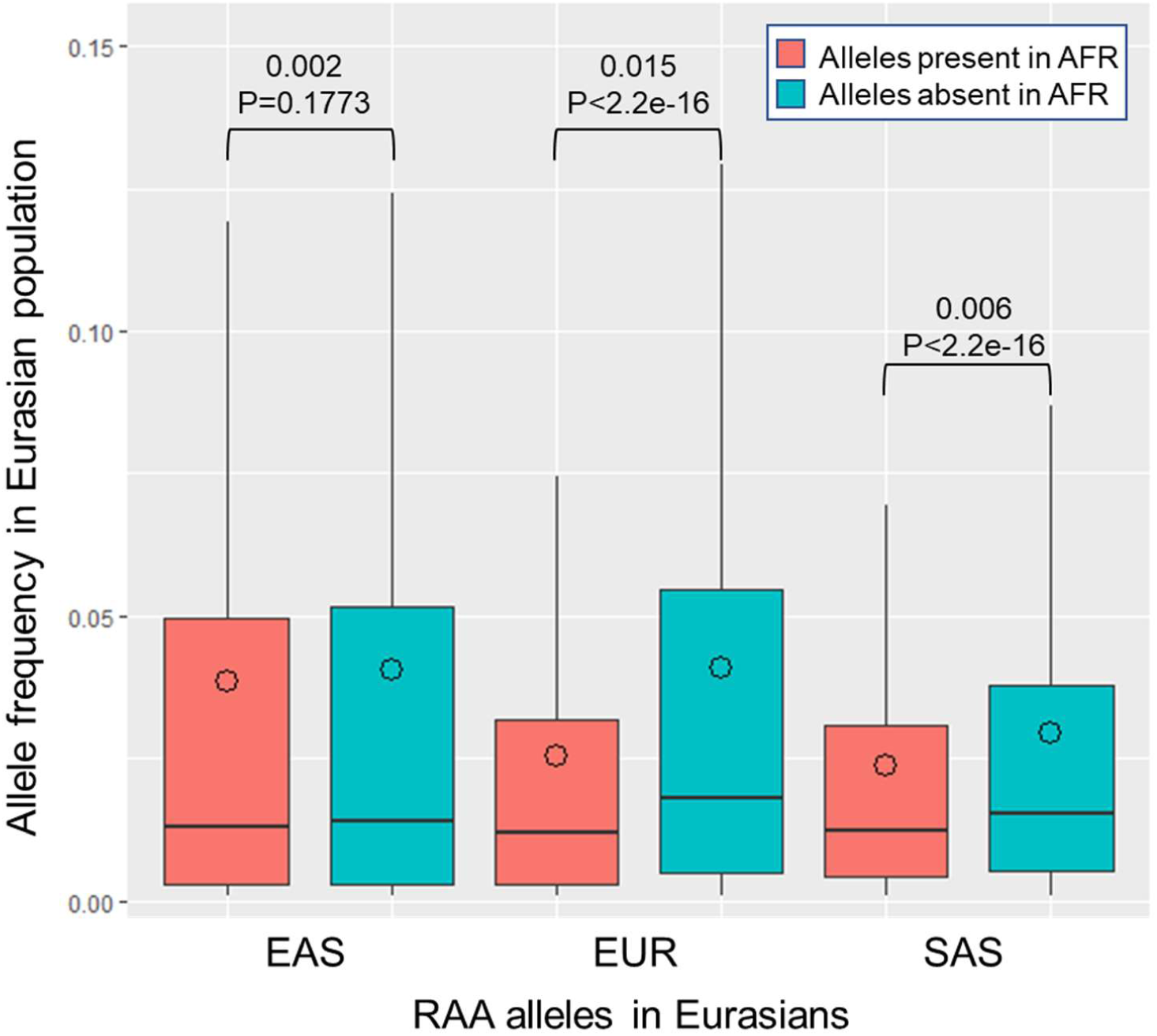
Comparison of allele frequencies across three Eurasian populations stratified by presence/absence of allele in modern sub-Saharan African populations. Reintroduced Ancestral Alleles (RAAs) that are also present in modern African (AFR) populations segregate at higher allele frequencies (AF) in all Eurasian populations than RAAs for which the allele is absent in AFR. Intra-population median differences in AF are displayed along with *P*-values (Mann Whitney U test). Outliers are not shown. Circles indicate mean AF.

**Figure S7.**
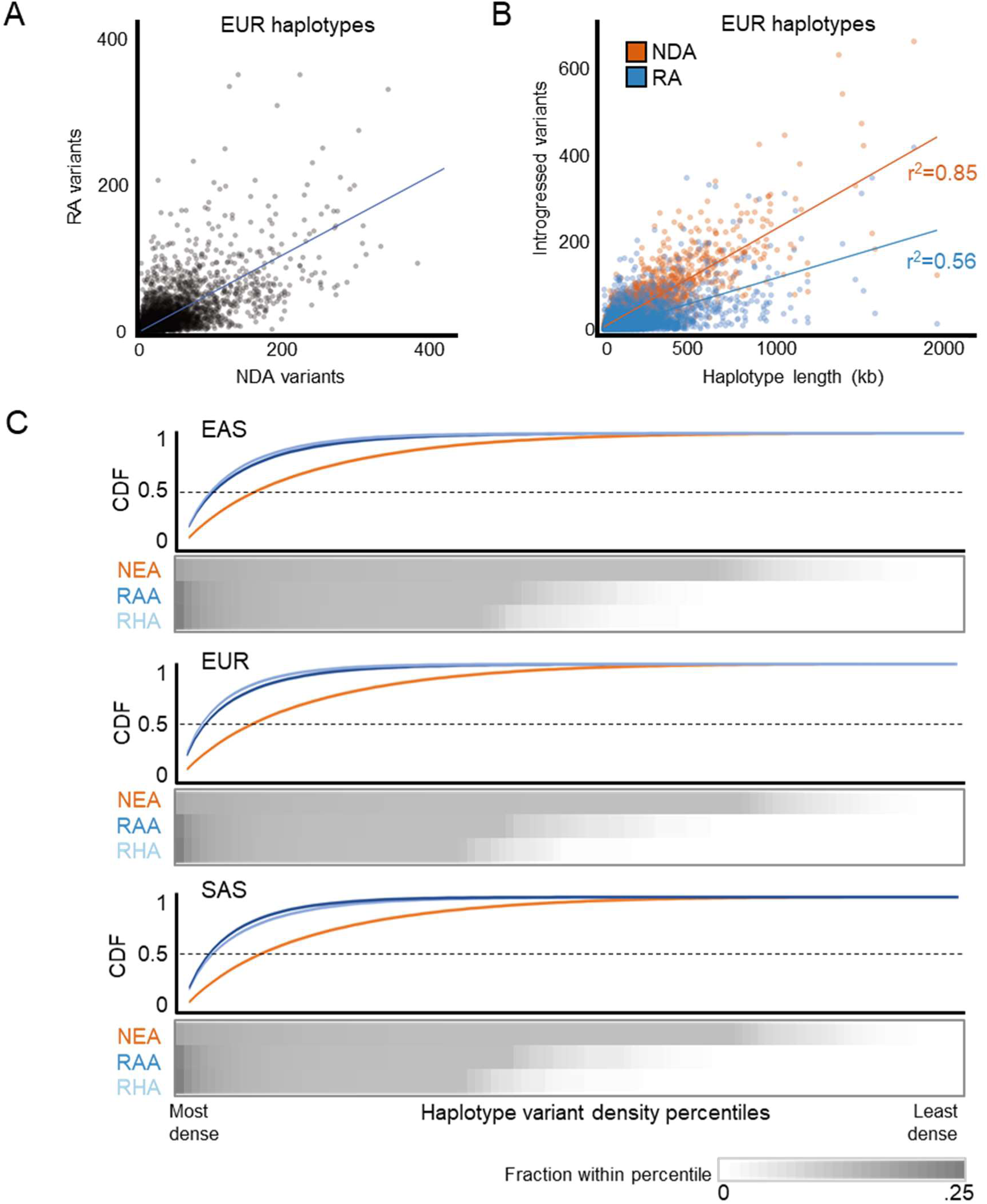
Reintroduced alleles cluster within introgressed Neanderthal haplotypes. (A) Scatter plot of the numbers of RAs and NDAs contained on all introgressed haplotypes in EUR. The correlation between the NDA and RA content is moderate (Pearson’s r^2^=0.46), with 18% of the haplotypes containing no RAs and 10% having more RAs than NDAs. (B) Scatter plot of the number of introgressed variants on each haplotype vs. haplotype length. The NDA content of a haplotype is proportional to its length (r^2^ = 0.85), but the number of RAs in each haplotype is less strongly correlated with length (r^2^ = 0.56). (C) Heatmap of the fraction of NDAs and RAs in density percentiles (high to low, left to right) averaged over all introgressed Eurasian haplotypes. This information is summarized in a cumulative density function (CDF) above the heatmaps. A higher fraction of all RAs are found in the most dense percentiles; this reflects the fact that RAs are often present in more dense clusters than are NDAs.

**Figure S8.**
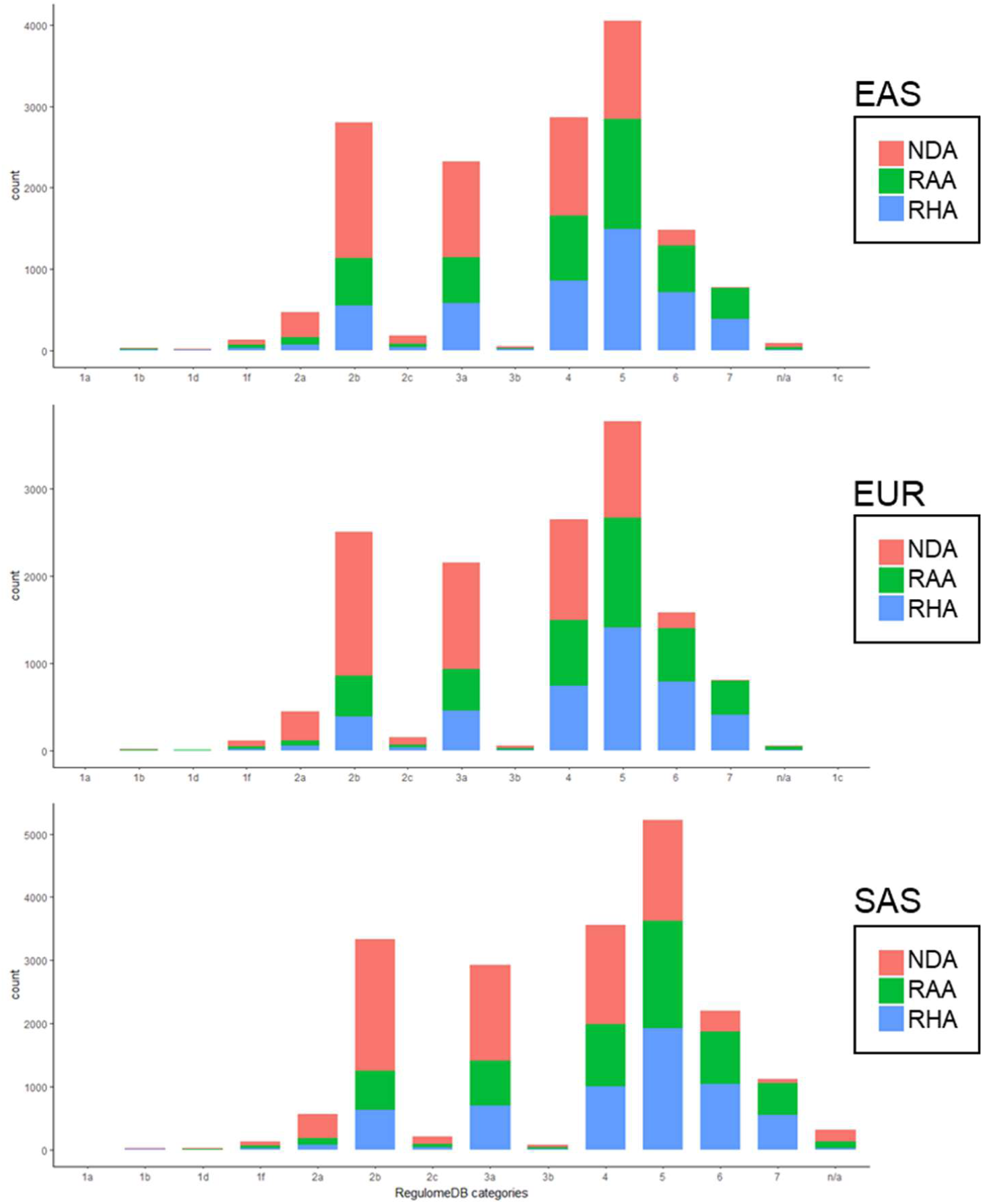
RAs and NDAs have similar amounts of overlap with annotated regulatory elements. Comparison of the fraction of NDAs and RAs in each of the RegulomeDB functional classes in order of evidence of regulatory activity.

**Figure S9.**
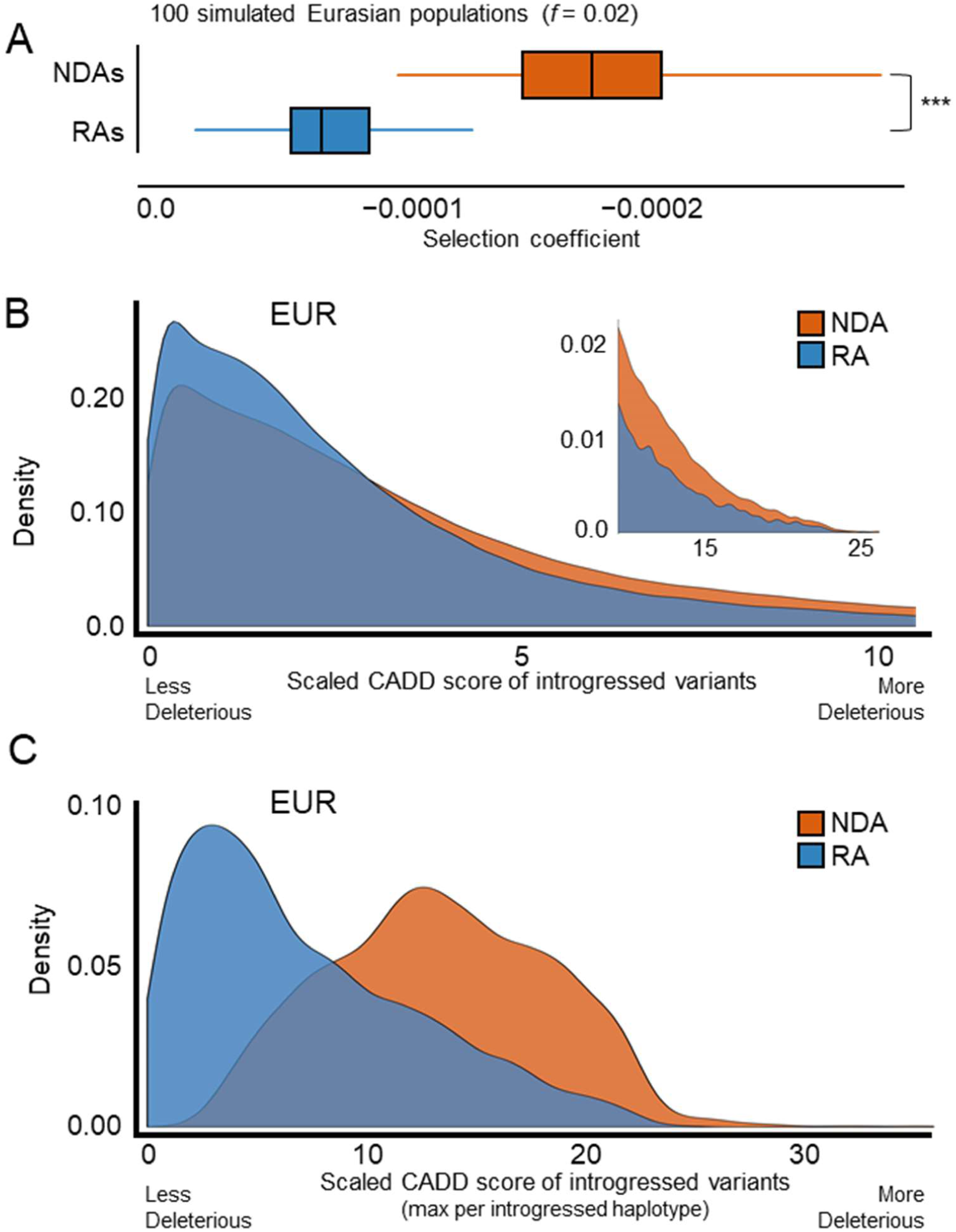
Reintroduced alleles have different predicted fitness effects than Neanderthal-derived alleles. (A) Simulations indicate that the RAs persisting in modern Eurasian populations are consistently less deleterious than NDAs over 200 simulations (median selection coefficient RA=7.7e-5; NDA=1.9e-4, P ≈ 0, Mann Whitney U test test). (B) In modern European (EUR) populations, RAs are predicted to be significantly less deleterious than NDAs by CADD (median scaled CADD: NDA=2.7; RA=2.1; P ≈ 0). The upper tail of highly deleterious mutations is highlighted in the inset. Results are similar for unscaled scores. (C) At the haplotype level, the maximum RA CADD score per introgressed haplotype is significantly lower than for NDAs (median scaled max CADD: NDA=13.3; RA=5.8; P ≈ 0). This is in part due to the overall difference demonstrated in (B) and to the greater number of NDAs per haplotype. RAs are rarely the most deleterious variant per haplotype. Results in East Asian and South Asian populations are similar (**Figure** S12).

**Figure S10.**
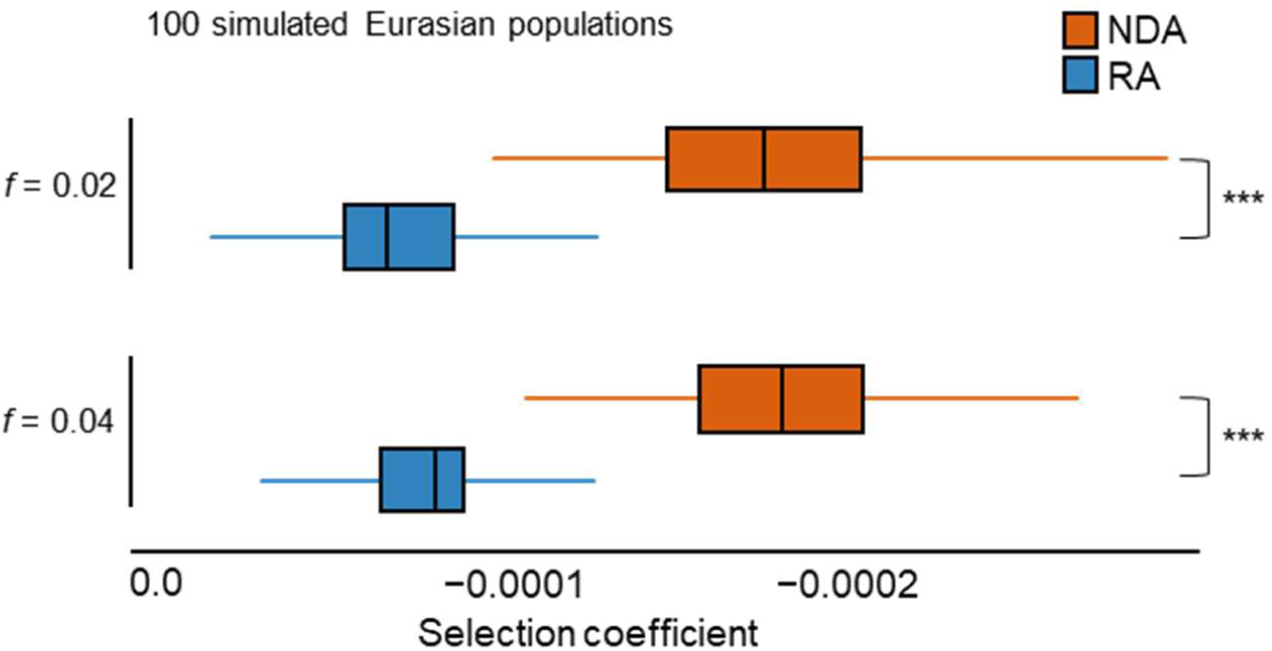
Distribution of selection coefficients of introgressed variants in simulated modern Eurasian populations are similar between admixture fractions. Selection coefficients in Eurasians from SLiM simulations with high (0.04) and low (0.02) admixture fractions. Each boxplot summarizes the average selection coefficient of all alleles in each introgressed class in each of 100 simulated modern Eurasian populations.

**Figure S11.**
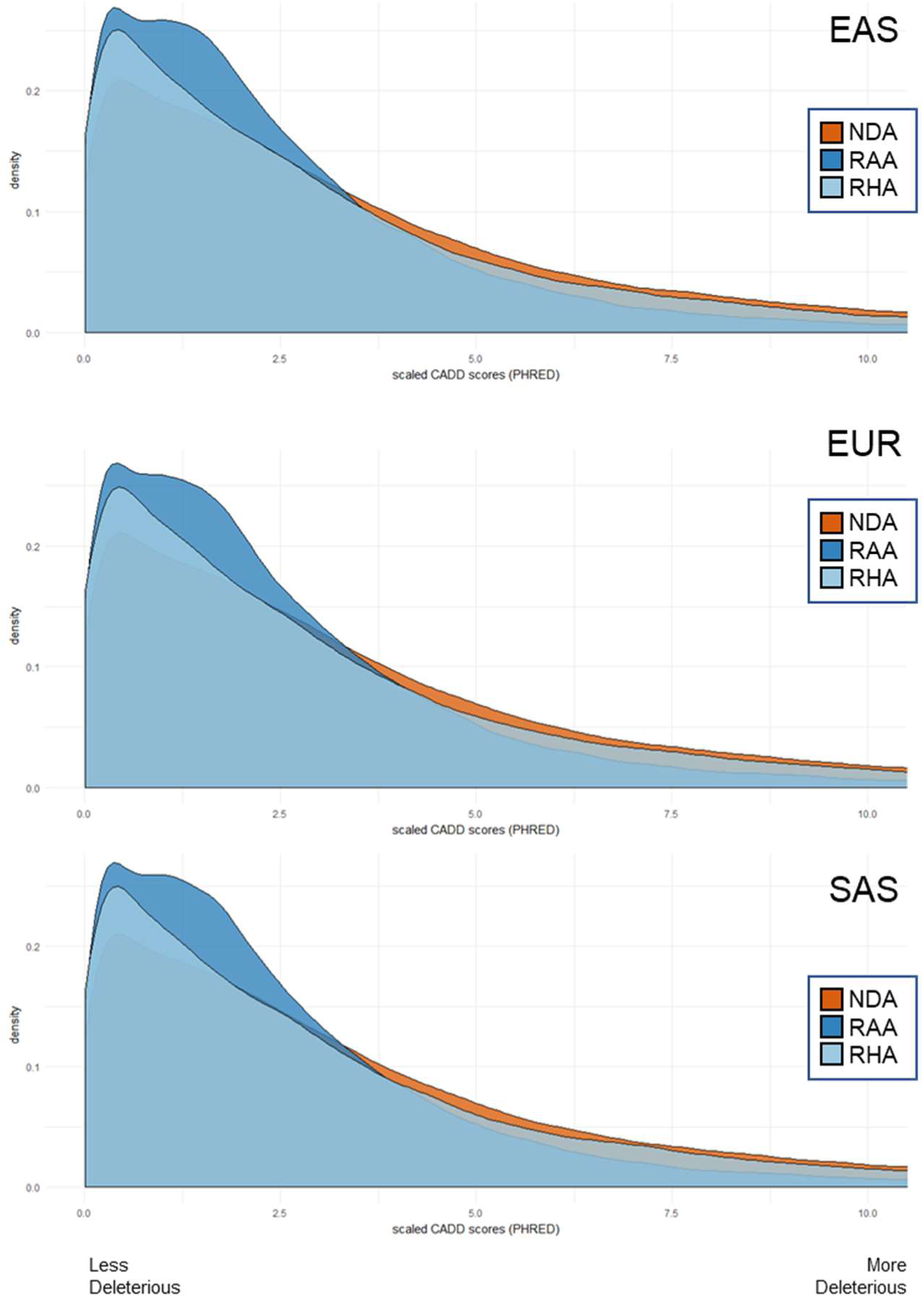
CADD scores for RAs (stratified as RAA and RHA) and NDAs in each of three populations. Normalized CADD scores for the introgressed variant classes (RAs and NDAs) with RAs separated into RAAs and RHAs. Considering RAAs and RHAs separately revealed that the RAAs are less deleterious than the RHAs (median scaled CADD score: RAA=1.91; RHA=2.23; P = 1.80e-89, Mann Whitney U test). This difference likely reflects the greater evolutionary conservation of RAAs. Results were similar across each superpopulation

**Figure S12.**
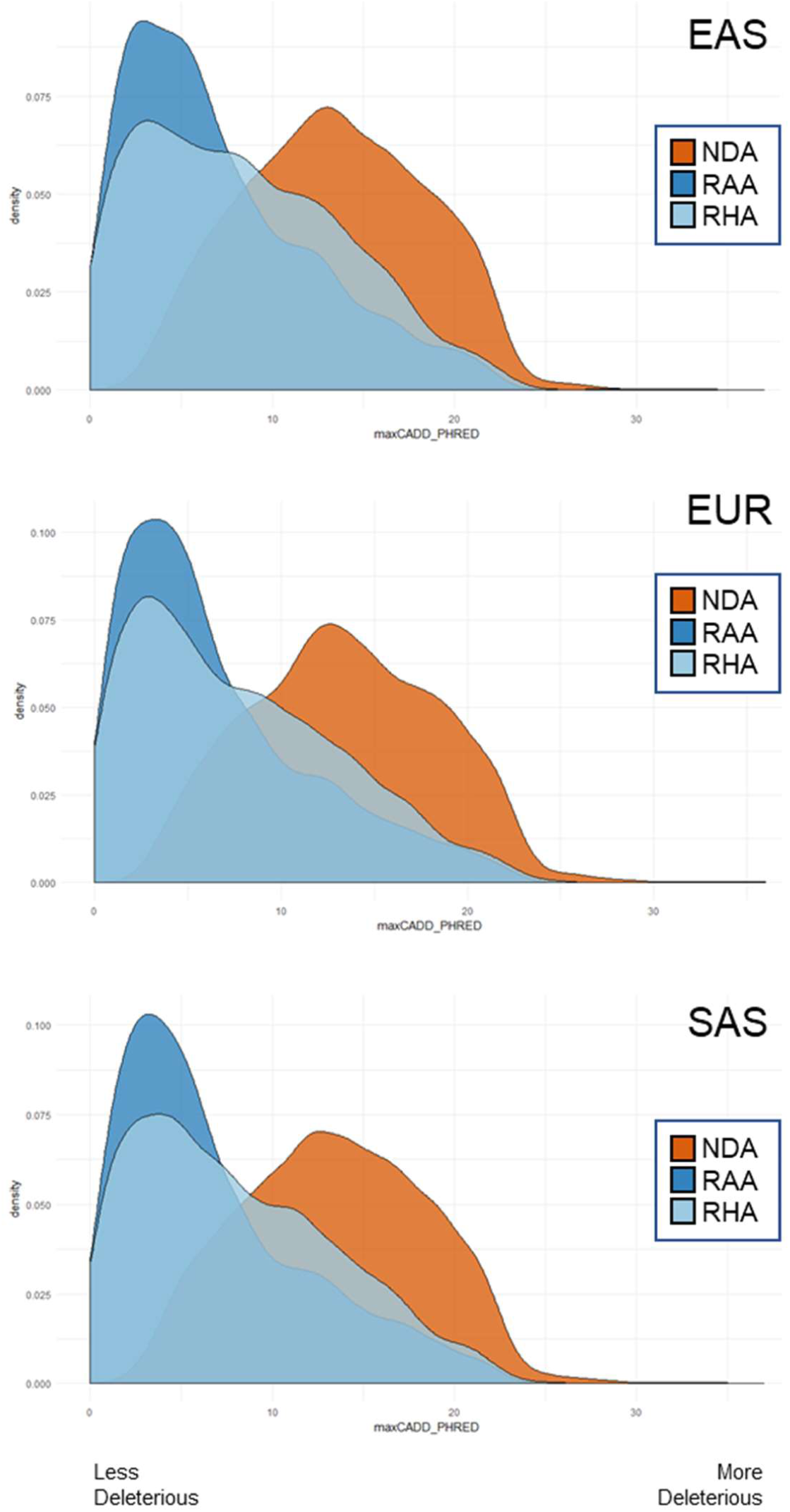
Max scaled CADD score per introgressed haplotype. The maximum scaled CADD score for each class of introgressed variant across introgressed haplotypes in each of three Eurasian populations. Maximum CADD scores of each class of RA (RAA and RHA) are much lower than those of the NDAs (P = 0, Mann Whitney U test).

**Figure S13.**
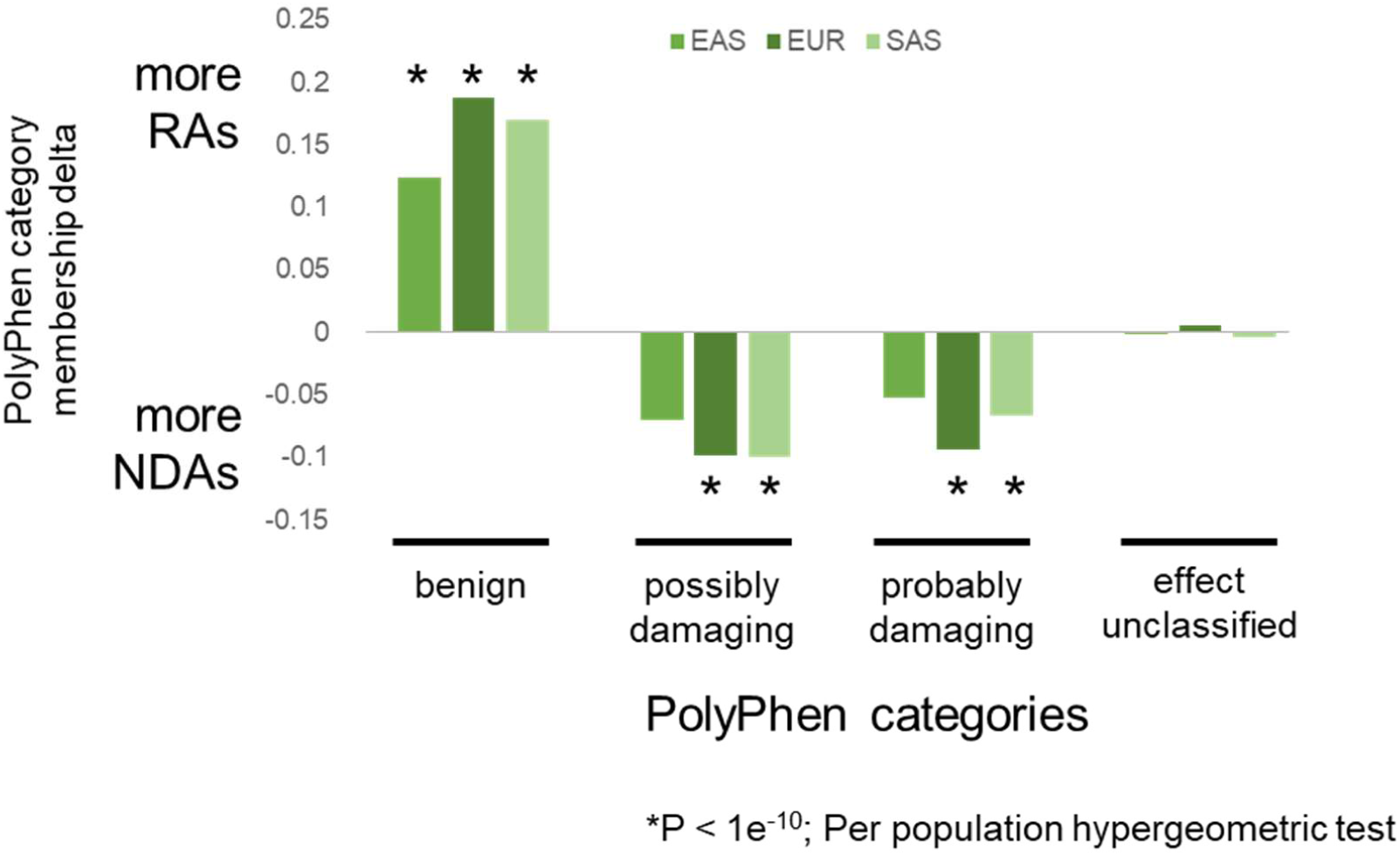
PolyPhen2 predicts RAs to be less damaging than NDAs. PolyPhen2 is more likely to classify RAs as “benign” in all three Eurasian populations. Conversely, NDAs are significantly more likely to be classified as “damaging” in both EUR and SAS populations. The y-axis reports the difference in PolyPhen category membership for each population (i.e., the fraction all RAs in population in the PolyPhen category minus the fraction of all NDAs in population in the PolyPhen category). Per population hypergeometric test is calculated on the enrichment (positive delta) or depletion (negative delta) for RA content within each PolyPhen category.

**Figure S14.**
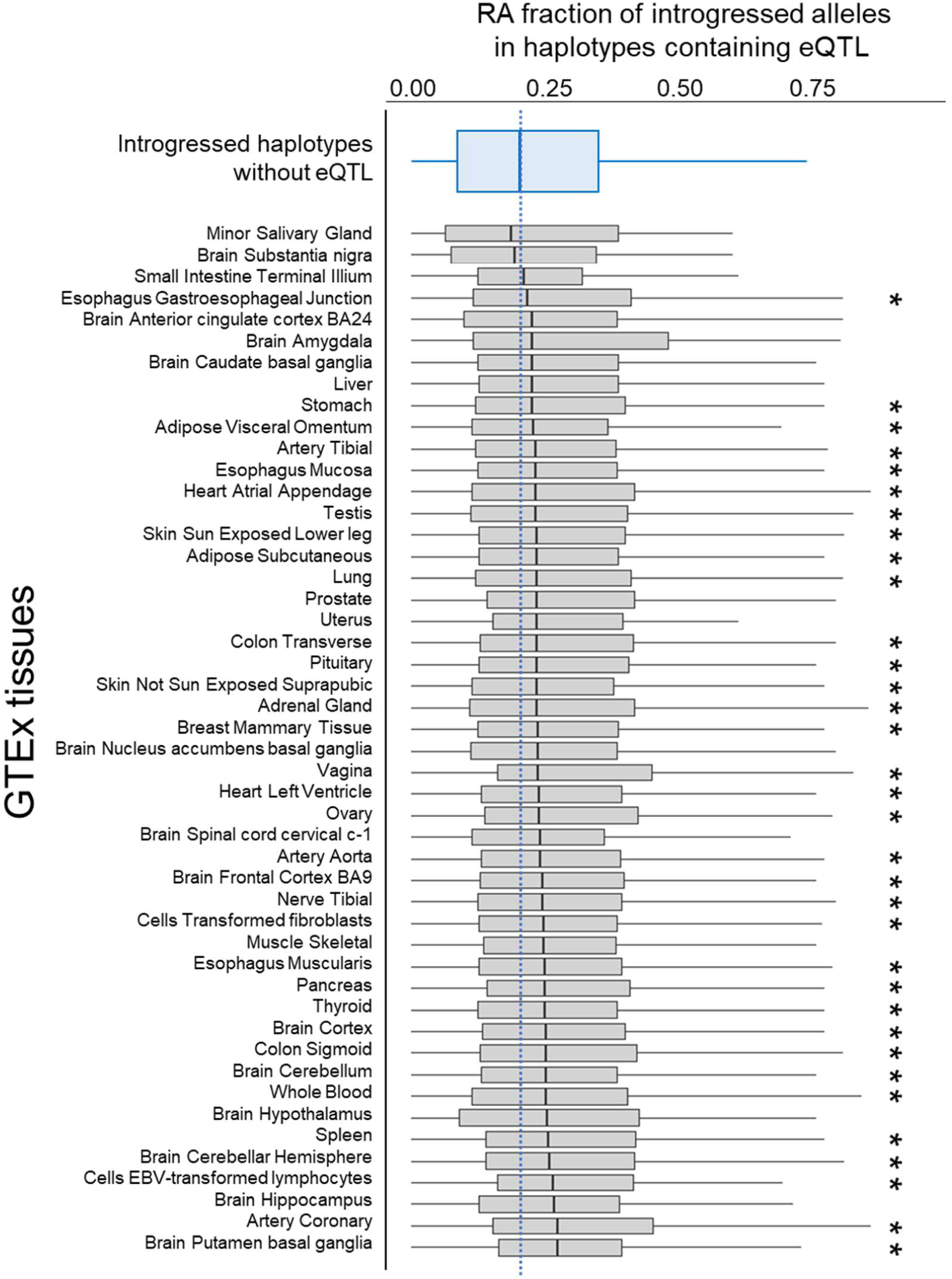
RA fraction in introgressed haplotypes containing eQTL in GTEx tissues. Summary of the RA fraction among introgressed variants in Neanderthal haplotypes in Europeans (EUR). Boxplots show the distributions RA fractions of all haplotypes containing at least one introgressed eQTL (RA or NDA) in the given GTEx tissue (gray box plots). These distributions are then compared pairwise with distribution for introgressed haplotypes that contain no introgressed GTEx eQTL (top, blue boxplot; n=4237). Haplotypes containing GTEx eQTL have RA contents higher than non-eQTL containing haplotypes in 46 tissues, with 34 of the tissues (*) having a significantly higher the RA fraction (P<0.05, Mann Whitney U Test).

## SUPPLEMENTARY TABLES

**Table S1.**
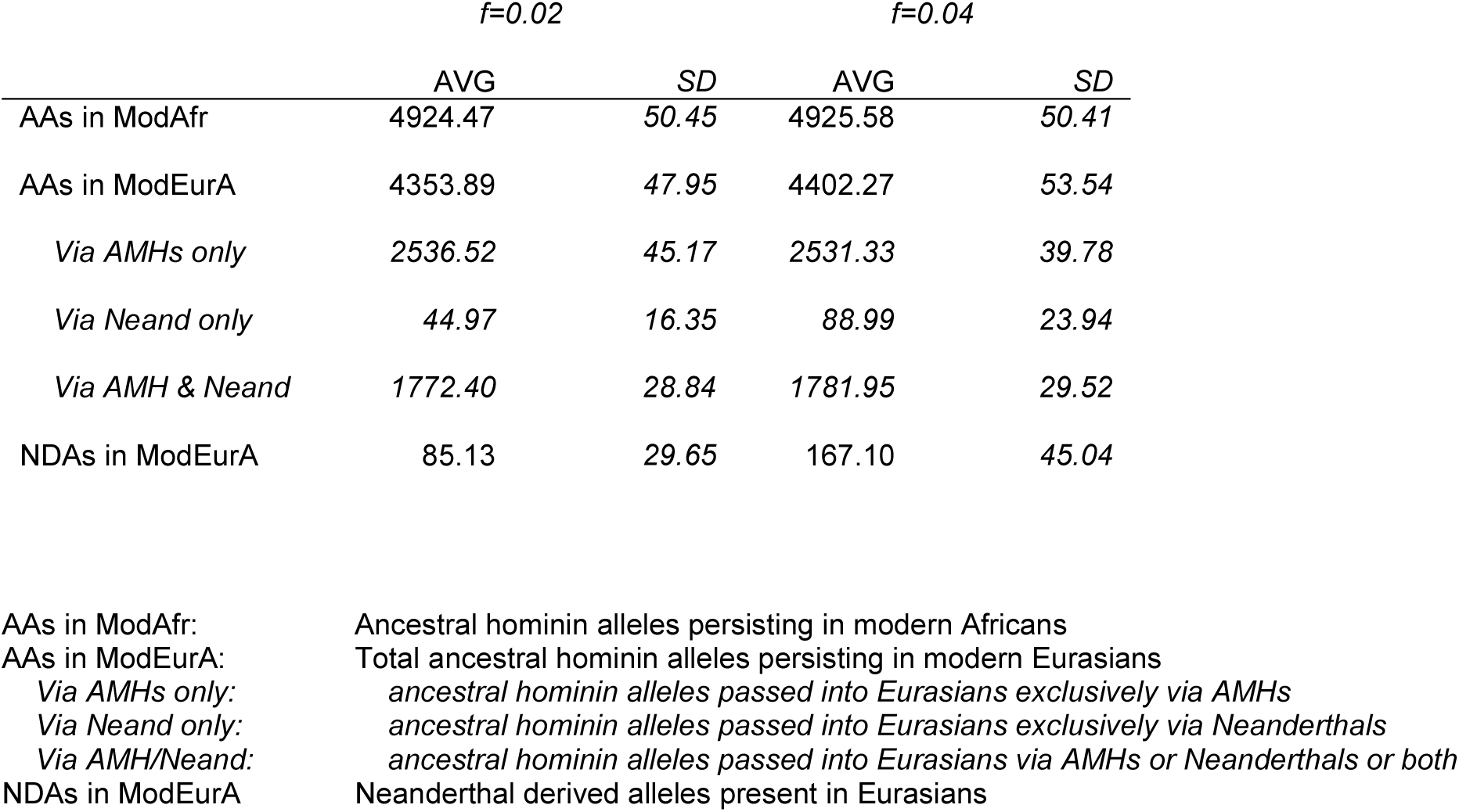
Counts of simulated ancient allele trajectories into modern Eurasians.

|  | *f*=0.02 |  | *f*=0.04 |  |
| --- | --- | --- | --- | --- |
|  | AVG | SD | AVG | SD |
| AAs in ModAfr | 4924.47 | 50.45 | 4925.58 | 50.41 |
| AAs in ModEurA | 4353.89 | 47.95 | 4402.27 | 53.54 |
| *Via AMHs only* | 2536.52 | 45.17 | 2531.33 | 39.78 |
| *Via Neand only* | 44.97 | 16.35 | 88.99 | 23.94 |
| *Via AMH & Neand* | 1772.40 | 28.84 | 1781.95 | 29.52 |
| NDAs in ModEurA | 85.13 | 29.65 | 167.10 | 45.04 |

|  |  |
| --- | --- |
| AAs in ModAfr: | Ancestral hominin alleles persisting in modern Africans |
| AAs in ModEurA: | Total ancestral hominin alleles persisting in modern Eurasians |
| *Via AMHs only:* | *ancestral hominin alleles passed into Eurasians exclusively via AMHs* |
| *Via Neand only:* | *ancestral hominin alleles passed into Eurasians exclusively via Neanderthals* |
| *Via AMH/Neand:* | *ancestral hominin alleles passed into Eurasians via AMHs or Neanderthals or both* |
| NDAs in ModEurA | Neanderthal derived alleles present in Eurasians |

**Table S2.** Simulations suggest that recombination of segregating human alleles onto introgressed haplotypes rarely results in false ascertainment of RAs.

|  | *f*=0.02 | *f*=0.04 |
| --- | --- | --- |
| False Discovery Rate (FP / (TP + FP)) | 0.009 | 0.0165 |

**Table S3.**
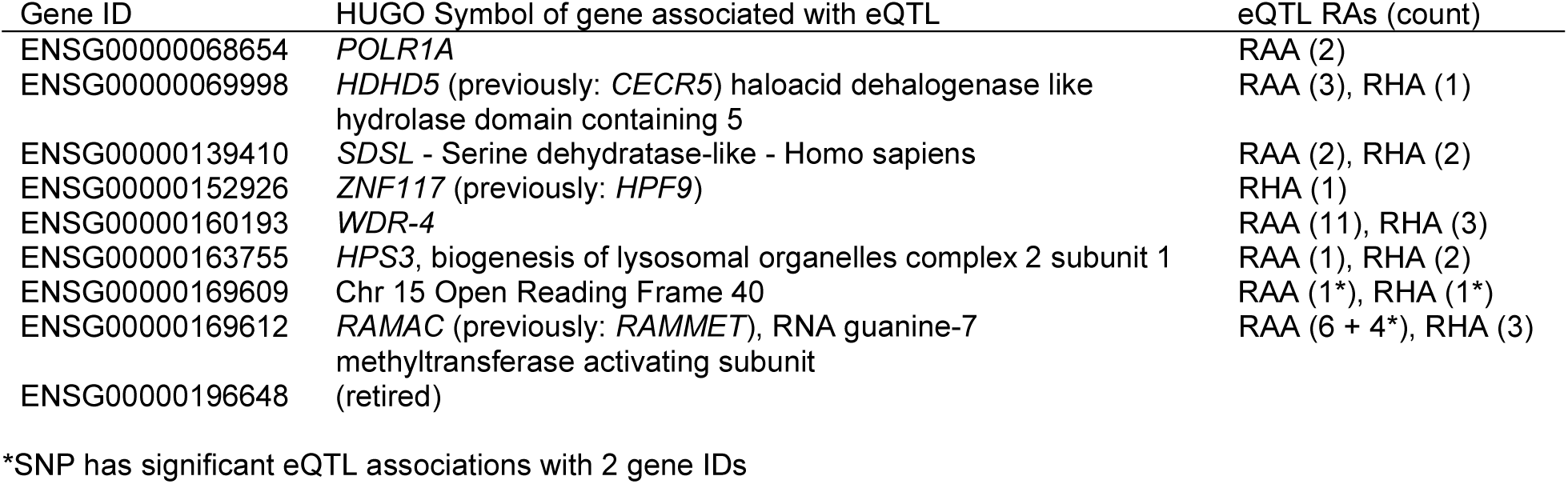
Genes associated with RA eQTL in LCLs shared between EUR and YRI.

| Gene ID | HUGO Symbol of gene associated with eQTL | eQTL RAs (count) |
| --- | --- | --- |
| ENSG00000068654 | *POLR1A* | RAA (2) |
| ENSG00000069998 | *HDHD5* (previously: *CECR5*) haloacid dehalogenase like hydrolase domain containing 5 | RAA (3), RHA (1) |
| ENSG00000139410 | *SDSL* - Serine dehydratase-like - Homo sapiens | RAA (2), RHA (2) |
| ENSG00000152926 | *ZNF117* (previously: *HPF9*) | RHA (1) |
| ENSG00000160193 | *WDR-4* | RAA (11), RHA (3) |
| ENSG00000163755 | *HPS3*, biogenesis of lysosomal organelles complex 2 subunit 1 | RAA (1), RHA (2) |
| ENSG00000169609 | Chr 15 Open Reading Frame 40 | RAA (1*), RHA (1*) |
| ENSG00000169612 | *RAMAC* (previously: *RAMMET*), RNA guanine-7 methyltransferase activating subunit | RAA (6 + 4*), RHA (3) |
| ENSG00000196648 | (retired) |  |

\*SNP has significant eQTL associations with 2 gene IDs

**Table S4.** European introgressed variants with significant MPRA evidence of modulating expression in LCLs.

| Chr | Loc (hg19) | Introgressed class | rsID | nearest TSS | distance from TSS | HUGO name |
| --- | --- | --- | --- | --- | --- | --- |
| chr2 | 68490970 | NDA | rs12713637 | NM_000945 | 11318 | *PPP3R1* |
| chr2 | 121044922 | NDA | rs79742700 | NM_002881 | 34508 | *RALB* |
| chr3 | 119188213 | NDA | rs75416321 | NM_152305 | 428 | *POGLUT1* |
| chr4 | 38805942 | NDA | rs5743566 | NM_003263 | 470 | *TLR1* |
| chr4 | 38807328 | NDA | rs45588337 | NM_003263 | 915 | *TLR1* |
| chr7 | 64448291 | RHA | rs11972247 | NM_015852 | 3123 | *ZNF117* |
| chr9 | 101969304 | RAA | rs77521170 | NM_033087 | 14942 | *ALG2* |
| chr12 | 113364471 | NDA | rs4767034 | NM_006187 | 11777 | *OAS3* |
| chr12 | 132569521 | RHA | rs77330556 | NR_003290 | 693 | *EP400NL* |
| chr19 | 41209256 | RAA | rs116796128 | NM_001142555 | 11755 | *ADCK4* |
| chr22 | 17639918 | RAA | rs71312076 | NM_033070 | 251 | *HDHD5* |

**Table S5.**
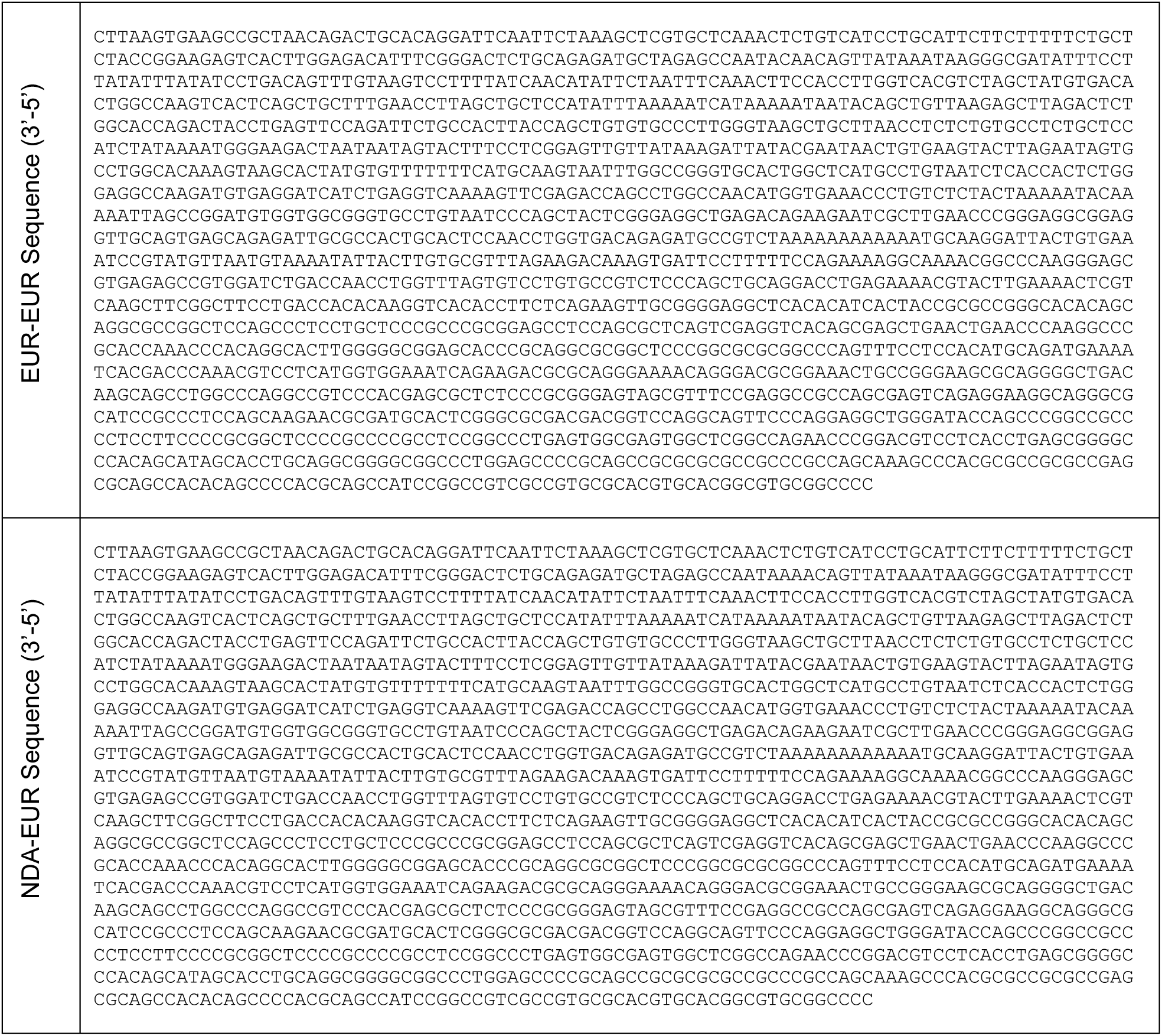
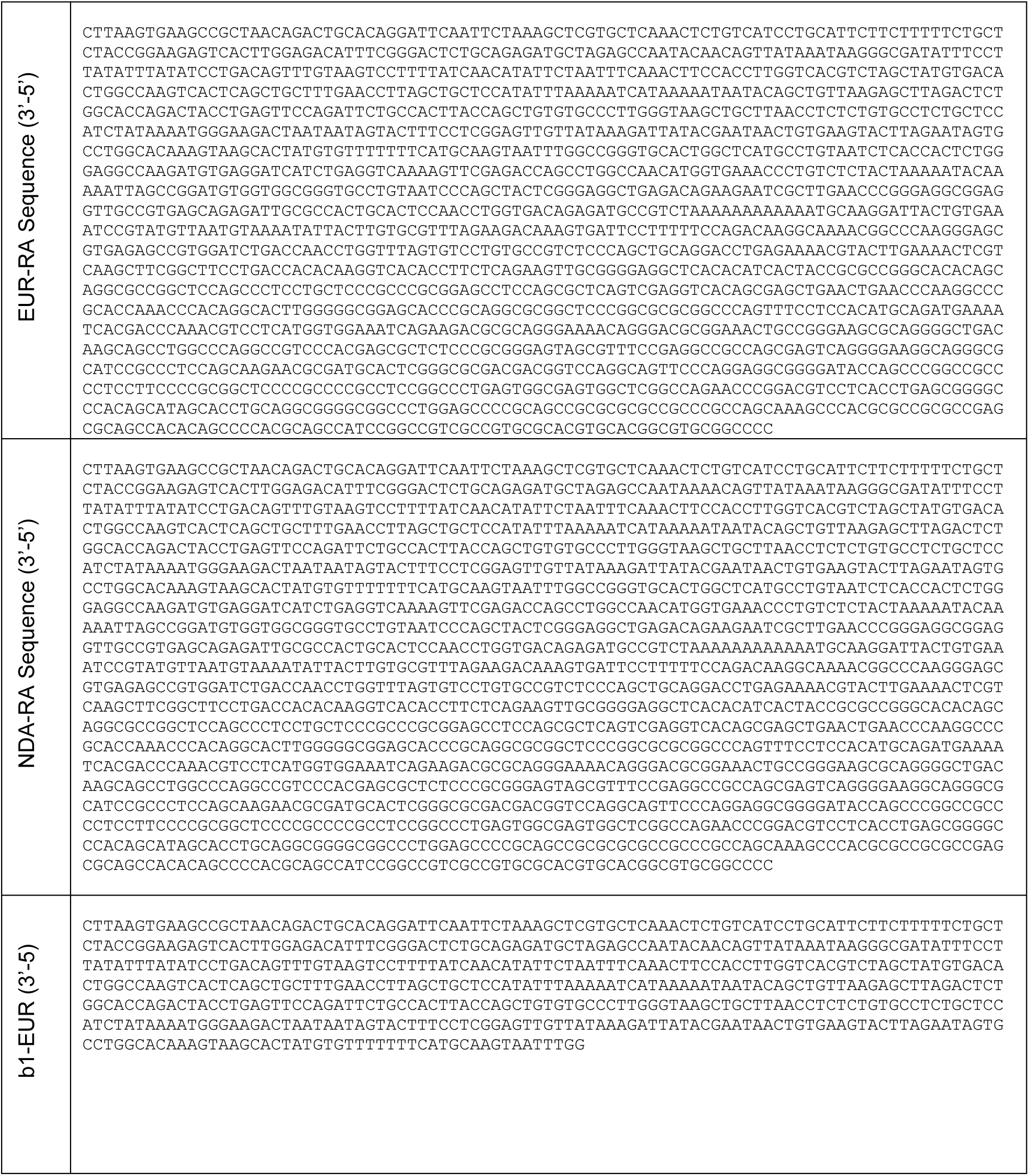
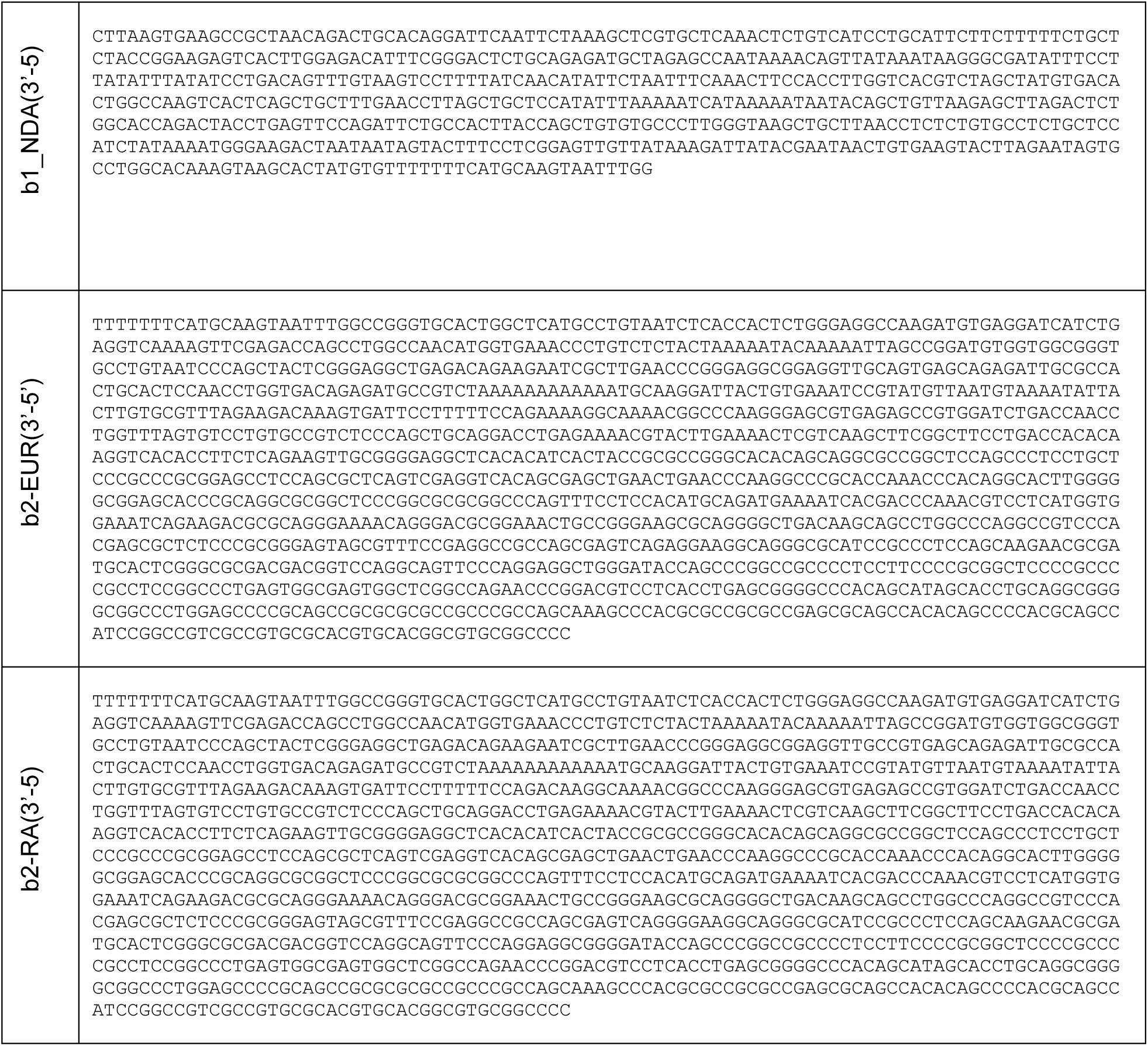
Luciferase reporter insert sequences.

**Table S6.**
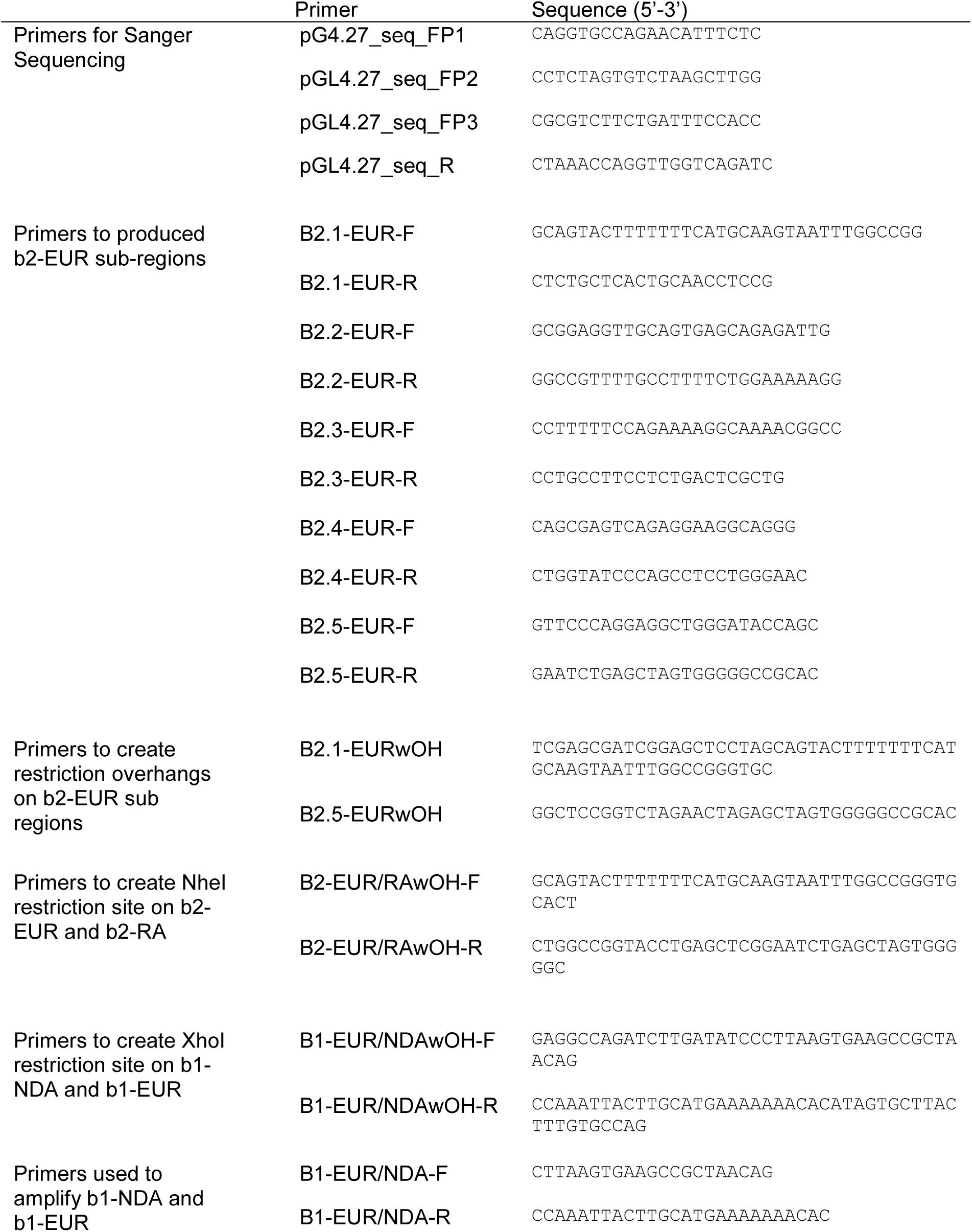
Luciferase primer sequences.

**Table S7.**
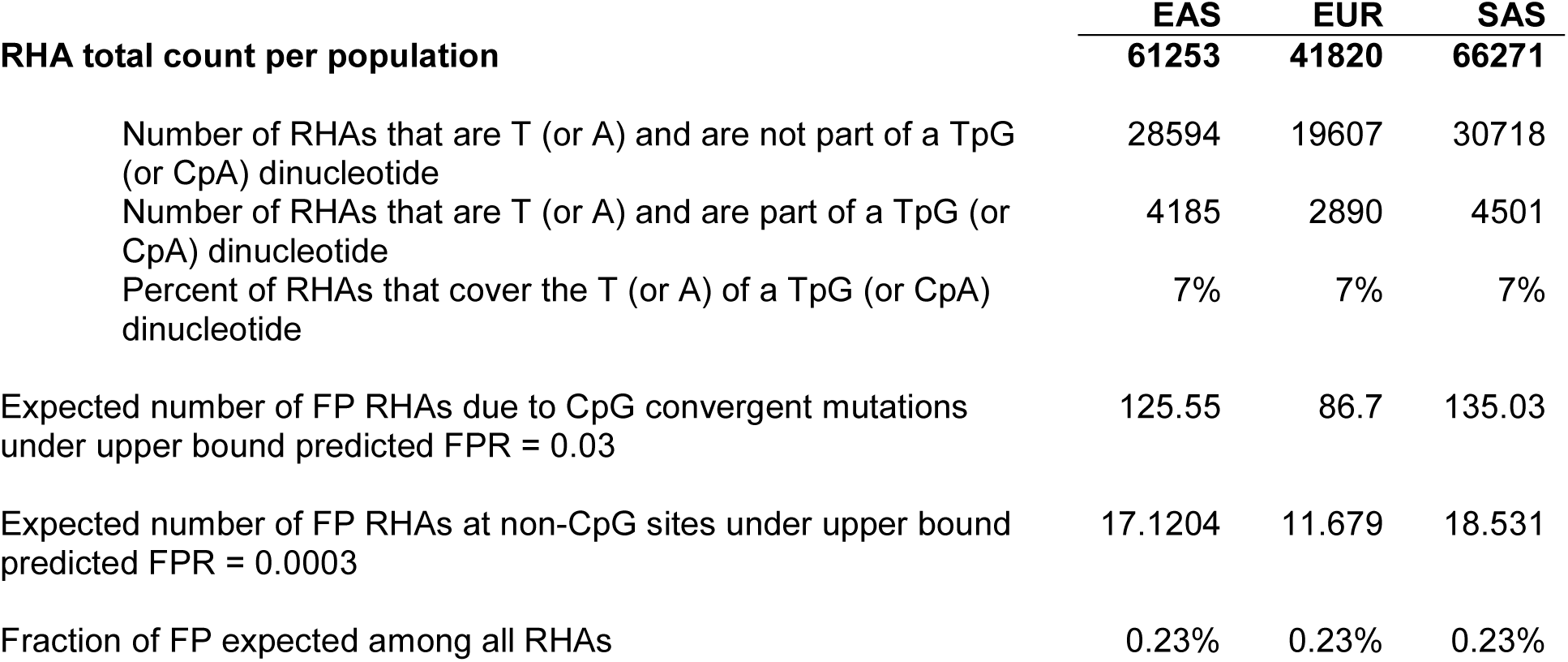
Occurrence of RHAs at putatively ancestral CpG.

## SUPPLEMENTARY FILE LIST

**File S1 List of NDAs and RAs for each Eurasian population**

**File S2 Table of GWAS hits for NDAs and RAs**

## REFERENCES

1. The 1000 Genomes Project Consortium, A global reference for human genetic variation. Nature. 526, 68–74 (2015).

2. S. Mallick et al., The Simons genome diversity project: 300 genomes from 142 diverse populations. Nature. 538, 201 (2016).

3. L. Pagani et al., Genomic analyses inform on migration events during the peopling of Eurasia. Nature. 538, 238–242 (2016).

4. B. M. Henn, L. R. Botigué, C. D. Bustamante, A. G. Clark, S. Gravel, Estimating Mutation Load in Human Genomes. Nat. Rev. Genet. 16, 333–343 (2015).

5. K. Prüfer et al., The complete genome sequence of a Neanderthal from the Altai Mountains. Nature. 505, 43–49 (2014).

6. K. Prüfer et al., A high-coverage Neandertal genome from Vindija Cave in Croatia. Science (80-.). 358, 655–658 (2017).

7. M. Meyer et al., A high-coverage genome sequence from an archaic Denisovan individual. Science (80-.). 338, 222–226 (2012).

8. R. E. Green et al., A draft sequence of the neandertal genome. Science (80-.). 328, 710–722 (2010).

9. S. Sankararaman et al., The genomic landscape of Neanderthal ancestry in present-day humans. Nature. 507, 354–357 (2014).

10. B. Vernot, J. M. Akey, Science (80-.)., in press.

11. S. Sankararaman, S. Mallick, N. Patterson, D. Reich, The Combined Landscape of Denisovan and Neanderthal Ancestry in Present-Day Humans. Curr. Biol. 26, 1241–1247 (2016).

12. B. Vernot et al., Excavating Neandertal and Denisovan DNA from the genomes of Melanesian individuals. Science (80-.). 352, 235–239 (2016).

13. L. Abi-Rached et al., The Shaping of Modern Human Immune Systems by Multiregional Admixture with Archaic Humans. Science (80-.). 334, 89–94 (2011).

14. F. L. Mendez, J. C. Watkins, M. F. Hammer, A Haplotype at STAT2 Introgressed from Neanderthals and Serves as a Candidate of Positive Selection in Papua New Guinea. Am. J. Hum. Genet. 91, 265–274 (2012).

15. M. Dannemann, K. Prüfer, J. Kelso, Functional implications of Neandertal introgression in modern humans. Genome Biol. 18, 61 (2017).

16. X. Liu, X. Jian, E. Boerwinkle, dbNSFP: a lightweight database of human nonsynonymous SNPs and their functional predictions. Hum. Mutat. 32, 894–9 (2011).

17. F. Racimo, S. Sankararaman, R. Nielsen, E. Huerta-Sánchez, Evidence for archaic adaptive introgression in humans. Nat. Rev. Genet. 16, 359–371 (2015).

18. F. Racimo, D. Marnetto, E. Huerta-Sánchez, Signatures of archaic adaptive introgression in present-day human populations. Mol. Biol. Evol. (2017), doi:10.1093/molbev/msw216.

19. I. Juric, S. Aeschbacher, G. Coop, The Strength of Selection against Neanderthal Introgression. PLOS Genet. 12, e1006340 (2016).

20. K. Harris, R. Nielsen, The Genetic Cost of Neanderthal Introgression. Genetics (2016).

21. R. M. Gittelman et al., Archaic Hominin Admixture Facilitated Adaptation to Out-of-Africa Environments. Curr. Biol. 26, 3375–3382 (2016).

22. M. Petr, S. Pääbo, J. Kelso, B. Vernot, The limits of long-term selection against Neandertal introgression. bioRxiv Prepr., 362566 (2018).

23. M. Dannemann, J. Kelso, The Contribution of Neanderthals to Phenotypic Variation in Modern Humans. Am. J. Hum. Genet. 101, 578–589 (2017).

24. C. N. Simonti et al., The phenotypic legacy of admixture between modern humans and Neandertals. Science (80-.). 351, 737–741 (2016).

25. Y. Nédélec et al., Genetic Ancestry and Natural Selection Drive Population Differences in Immune Responses to Pathogens. Cell (2016), doi:10.1016/j.cell.2016.09.025.

26. H. Quach et al., Genetic Adaptation and Neandertal Admixture Shaped the Immune System of Human Populations. Cell (2016), doi:10.1016/j.cell.2016.09.024.

27. A. J. Sams et al., Adaptively introgressed Neandertal haplotype at the OAS locus functionally impacts innate immune responses in humans. Genome Biol. 17, 246 (2016).

28. Y. Hu, Q. Ding, Y. He, S. Xu, L. Jin, Reintroduction of a Homocysteine Level-Associated Allele into East Asians by Neanderthal Introgression. Mol. Biol. Evol. 32, msv176 (2015).

29. A. Bergström et al., Insights into human genetic variation and population history from 929 diverse genomes. bioRxiv, 674986 (2019).

30. F. A. Villanea, J. G. Schraiber, Multiple episodes of interbreeding between Neanderthal and modern humans. Nat. Ecol. Evol. 3, 39–44 (2019).

31. J. D. Wall et al., Higher Levels of Neanderthal Ancestry in East Asians than in Europeans. Genetics. 194, 199–209 (2013).

32. B. Y. Kim, K. E. Lohmueller, Selection and Reduced Population Size Cannot Explain Higher Amounts of Neandertal Ancestry in East Asian than in European Human Populations. Am. J. Hum. Genet. 96, 454–461 (2015).

33. J. MacArthur et al., The new NHGRI-EBI Catalog of published genome-wide association studies (GWAS Catalog). Nucleic Acids Res. 45, D896–D901 (2017).

34. A. Franke et al., Genome-wide meta-analysis increases to 71 the number of confirmed Crohn’s disease susceptibility loci. Nat. Genet. 42, 1118–25 (2010).

35. L. Jostins et al., Host-microbe interactions have shaped the genetic architecture of inflammatory bowel disease. Nature. 491, 119–24 (2012).

36. M. K. Lee et al., Genome-wide association study of facial morphology reveals novel associations with FREM1 and PARK2. PLoS One. 12, e0176566 (2017).

37. S. L. Park et al., Mercapturic Acids Derived from the Toxicants Acrolein and Crotonaldehyde in the Urine of Cigarette Smokers from Five Ethnic Groups with Differing Risks for Lung Cancer. PLoS One. 10, e0124841 (2015).

38. J. Spada et al., Genome-wide association analysis of actigraphic sleep phenotypes in the LIFE Adult Study. J. Sleep Res. 25, 690–701 (2016).

39. A. M. Kulminski et al., Strong impact of natural-selection-free heterogeneity in genetics of age-related phenotypes. Aging (Albany. NY). 10, 492–514 (2018).

40. S. M. Lutz et al., A genome-wide association study identifies risk loci for spirometric measures among smokers of European and African ancestry. BMC Genet. 16, 138 (2015).

41. GTEx Consortium, Genetic effects on gene expression across human tissues. Nature. 550, 204–213 (2017).

42. R. C. McCoy, J. Wakefield, J. M. Akey, Impacts of Neanderthal-Introgressed Sequences on the Landscape of Human Gene Expression. Cell. 168, 916–927.e12 (2017).

43. C. N. Simonti et al., The phenotypic legacy of admixture between modern humans and Neandertals. Science (80-.). 351, 737–741 (2016).

44. T. Lappalainen et al., Transcriptome and genome sequencing uncovers functional variation in humans. Nature. 501, 506–11 (2013).

45. OMIM, CAT EYE SYNDROME; CES, (available at https://www.omim.org/entry/115470).

46. R. Tewhey et al., Direct Identification of Hundreds of Expression-Modulating Variants using a Multiplexed Reporter Assay. Cell. 165, 1519–1529 (2016).

47. P. Moorjani et al., The History of African Gene Flow into Southern Europeans, Levantines, and Jews. PLoS Genet. 7, e1001373 (2011).

48. A. Platt, J. Hey, Recent African gene flow responsible for excess of old rare genetic variation in Great Britain. bioRxiv (2017), doi:10.1101/190066.

49. D. Brawand et al., The evolution of gene expression levels in mammalian organs. Nature. 478, 343–348 (2011).

50. S. Castellano et al., Patterns of coding variation in the complete exomes of three Neandertals. Proc. Natl. Acad. Sci. 111, 6666–6671 (2014).

51. B. M. Henn et al., Distance from sub-Saharan Africa predicts mutational load in diverse human genomes. Proc. Natl. Acad. Sci. U. S. A. 113, E440–9 (2016).

52. K. E. Lohmueller, The distribution of deleterious genetic variation in human populations. Curr. Opin. Genet. Dev. 29, 139–146 (2014).

53. K. E. Lohmueller et al., Proportionally more deleterious genetic variation in European than in African populations. Nature. 451, 994–997 (2008).

54. Y. B. Simons, M. C. Turchin, J. K. Pritchard, G. Sella, The deleterious mutation load is insensitive to recent population history. Nat. Genet. 46, 220–4 (2014).

55. P. Danecek et al., The variant call format and VCFtools. Bioinformatics. 27, 2156–2158 (2011).

56. A. Hodgkinson, A. Eyre-Walker, Variation in the mutation rate across mammalian genomes. Nat. Rev. Genet. 12, 756–766 (2011).

57. M. Kircher et al., A general framework for estimating the relative pathogenicity of human genetic variants. Nat. Genet. 46, 310–315 (2014).

58. B. C. Haller, P. W. Messer, SLiM 2: Flexible, interactive forward genetic simulations. Mol. Biol. Evol. (2017), doi:10.1093/molbev/msw211.

59. A. Eyre-Walker, M. Woolfit, T. Phelps, The Distribution of Fitness Effects of New Deleterious Amino Acid Mutations in Humans. Genetics. 173, 891–900 (2006).

60. S. Gravel et al., Demographic history and rare allele sharing among human populations. Proc. Natl. Acad. Sci. 108, 11983–11988 (2011).

